# Near-infrared imaging in fission yeast by genetically encoded biosynthesis of phycocyanobilin

**DOI:** 10.1101/2021.05.19.444883

**Authors:** Keiichiro Sakai, Yohei Kondo, Hiroyoshi Fujioka, Mako Kamiya, Kazuhiro Aoki, Yuhei Goto

## Abstract

Near-infrared fluorescent protein (iRFP) is a bright and stable fluorescent protein with excitation and emission maxima at 690 nm and 713 nm, respectively. Unlike the other conventional fluorescent proteins such as GFP, iRFP requires biliverdin (BV) as a chromophore because iRFP originates from bacteriophytochrome. Here, we report that phycocyanobilin (PCB) functions as a brighter chromophore for iRFP than BV, and biosynthesis of PCB allows live-cell imaging with iRFP in the fission yeast *Schizosaccharomyces pombe*. We initially found that fission yeast cells did not produce BV, and therefore did not show any iRFP fluorescence. The brightness of iRFP attached to PCB was higher than that of iRFP attached to BV *in vitro* and in fission yeast. We introduced SynPCB, a previously reported PCB biosynthesis system, into fission yeast, resulting in the brightest iRFP fluorescence. To make iRFP readily available in fission yeast, we developed an endogenous gene tagging system with iRFP and all-in-one integration plasmids, which contain genes required for the SynPCB system and the iRFP-fused marker proteins. These tools not only enable the easy use of iRFP in fission yeast and the multiplexed live-cell imaging in fission yeast with a broader color palette, but also open the door to new opportunities for near-infrared fluorescence imaging in a wider range of living organisms.

## INTRODUCTION

Fluorescent proteins (FPs) have become indispensable to visualize the biological processes in living cells and tissues (Lambert 2019). Green fluorescent protein (GFP), the most widely used FP, has been intensively modified to improve the brightness and the photo-, thermo-, and pH-stabilities, and to change the excitation and emission spectrum. Use of a variety of fluorescent proteins with different excitation and emission spectra enables multiplexed fluorescence imaging to monitor multiple biological events simultaneously at high spatial and temporal resolution.

Near-infrared fluorescent proteins have been developed through the engineering of phytochromes, which are photosensory proteins of plants, bacteria, and fungi (Chernov et al. 2017), or allophycocyanin, which is a light-harvesting phycobiliprotein of cyanobacteria (Rodriguez et al. 2016). RpBphP2 from photosynthetic bacteria was engineered as an iRFP (later renamed iRFP713) by truncation and the saturation mutagenesis (Filonov et al. 2011). Since the initial report of iRFP, tremendous efforts have been devoted to developing near-infrared FPs with higher brightness, monomer formation, and longer wavelength (Shcherbakova and Verkhusha 2013; Shcherbakova et al. 2016, 2018; Matlashov et al. 2020; Stepanenko et al. 2016; Oliinyk et al. 2019; Kamper et al. 2018; Yu et al. 2014; Rodriguez et al. 2016; Yu et al. 2015; Rogers, Johnson, and Firnberg 2019; Filonov et al. 2011; Fushimi et al. 2019). Unlike the canonical fluorescent proteins derived from jellyfish or coral, phytochromes and allophycocyanin require a linear tetrapyrrole as a chromophore such as biliverdin IXα (BV), phycocyanobilin (PCB), or phytochromobilin (PΦB); bacteriophytochromes bind to BV, allophycocyanin and cyanobacterial phytochromes bind to PCB, and plantal phytochromes bind to PΦB. These photosensory proteins autocatalytically form a covalent bond with the chromophore (Fushimi and Narikawa 2021). These linear tetrapyrroles are produced from heme (Terry and Lagarias 1991; Beale 1993). Heme-oxygenase (HO) catalyzes oxidative cleavage of heme to generate BV with the help of ferredoxin (Fd), an electron donor, and ferredoxin-NADP+ reductase (Fnr) (Cornejo, Willows, and Beale 1998). In cyanobacteria, PCB is produced from BV through PcyA, Fd, and Fnr, while in plants, PΦB is synthesized from BV using HY2, Fd, and Fnr (Muramoto et al. 1999; Frankenberg et al. 2001; Kohchi et al. 2001). To exploit phytochromes that are required for PCB or PΦB in other organisms, our group and others have demonstrated reconstitution of BV, PCB, and PΦB synthesis in bacteria, mammalian cells, frog eggs, the budding yeast, *Pichia*, and fission yeast (Mukougawa et al. 2006; Gambetta and Lagarias 2001; Tooley, Cai, and Glazer 2001; Landgraf et al. 2001; K. Müller et al. 2013; Uda et al. 2017; Kyriakakis et al. 2018; Hochrein et al. 2017; Shin et al. 2014).

As the fluorescence of iRFP depends on chromophore formation, the BV concentration is of critical importance for imaging iRFP (Fig. 1A). Indeed, it has been reported that the addition of purified BV increases the fluorescence of iRFPs (Shemetov, Oliinyk, and Verkhusha 2017; Piatkevich et al. 2017). Alternatively, genetic modifications such as the overexpression of heme oxygenase-1 (HO1), which catalyzes heme to generate BV, and the knock out of biliverdin reductase A (BVRA), which degrades BV to generate bilirubin, improve the brightness of iRFP through the additional accumulation of BV (Shemetov, Oliinyk, and Verkhusha 2017; Kobachi et al. 2020). On the other hand, because *Caenorhabditis elegans* produces little or no BV (Ding et al. 2017), it is not possible to image biological processes in this nematode simply by introducing the iRFP gene. In the case of multicellular organisms that cannot produce BV, including *C*.*elegans*, the introduction of genes required for BV production is more effective than the external addition of BV because of the low tissue penetration property. However, at present, only the introduction of the HO1 gene has been reported as a genetically encoded method for inducing the iRFP chromophore, and it has not been improved or optimized yet.

**Fig 1.**
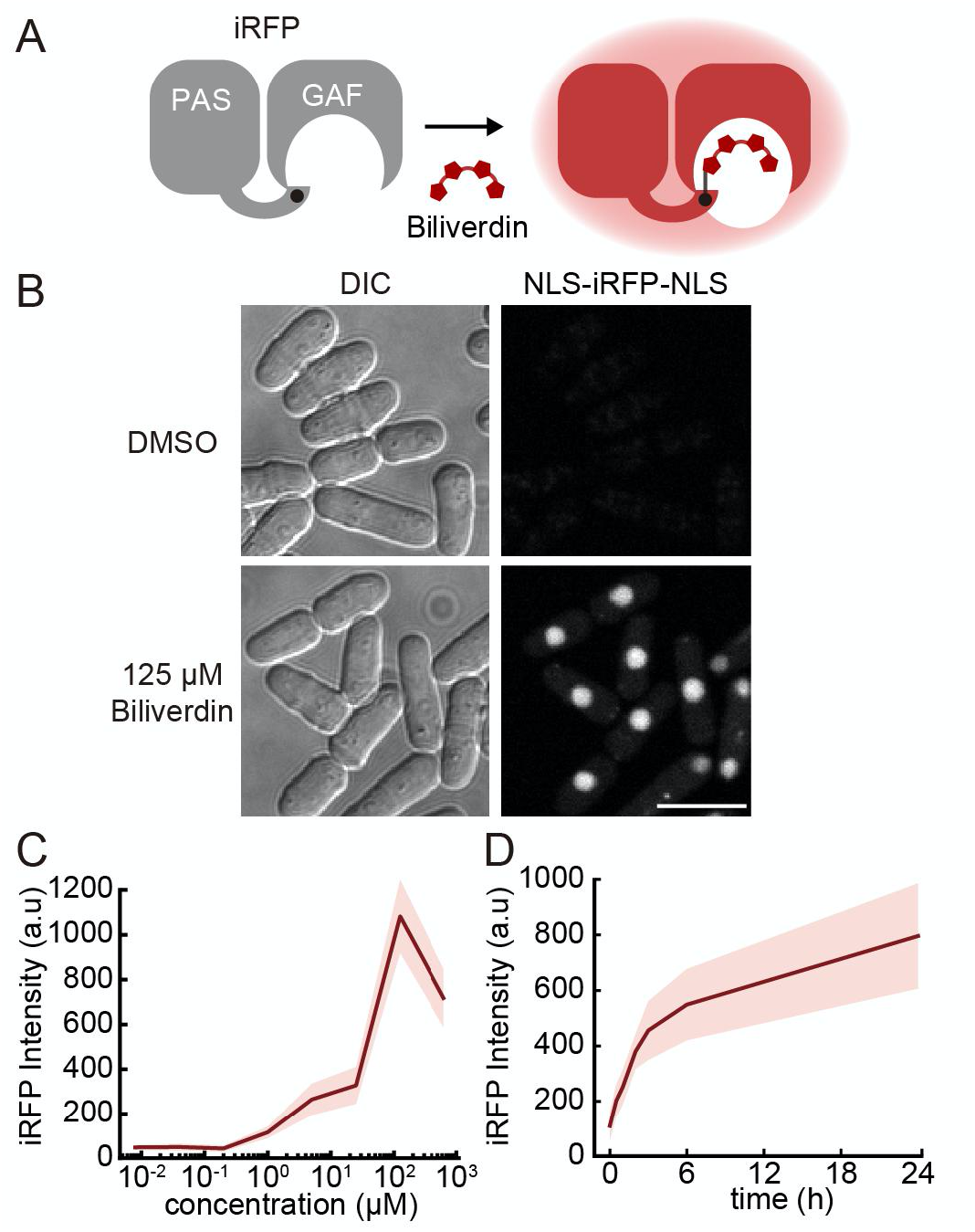
iRFP does not fluoresce in fission yeast. (A) Schematic illustration of chromophore formation of iRFP with biliverdin (BV). BV covalently attaches to iRFP as a chromophore. The PAS domain in iRFP contains a conserved cysteine residue at the N-terminus that covalently attaches to the BV, while the BV itself fits into the cleft in the GAF domain. (B) Representative images of fission yeast expressing NLS-iRFP-NLS with or without external BV treatment. Scale bar, 10 μm. (C) Dose-response curve of iRFP fluorescence as a function of levels of BV incorporation into fission yeast cells. Fission yeast cells were cultured in liquid YEA and incubated at room temperature for 3 h with the indicated concentration of BV (8 nM, 40 nM, 200 nM, 1 μM, 5 μM, 25 μM, 125 μM, and 625 μM). The red line and shaded area indicate the averaged intensity and S.D., respectively (n = 50 cells). The decrease in iRFP intensity under 625 μM BV could be due to cell death and/or toxicity by the excess DMSO. (D) Time-course of BV incorporation into fission yeast cells. Fission yeast cells were cultured in liquid YEA with the addition of 500 μM BV at 32°C with shaking, and cells were collected at the indicated points (0.5 h, 1 h, 2 h, 3 h, 6 h, 24 h). The red line and shaded area indicate the averaged intensity and S.D., respectively (n = 50 cells).

Here, we report that PCB acts as a better chromophore for iRFP than BV, and genetically encoded PCB synthesis outperforms HO1-mediated BV production in terms of iRFP brightness in fission yeast. We accidentally found that iRFP did not fluoresce in fission yeast because of the lack of the HO1 gene, and therefore the lack of BV. Both the external BV addition and heterologous HO1 expression rendered iRFP fluorescent in fission yeast. To our surprise, PCB biosynthesis with a SynPCB system, which we have previously reported (Uda et al. 2017, 2020), and treatment of the purified PCB yielded brighter iRFP fluorescence than that by either BV biosynthesis or BV treatment. We confirmed that PCB-bound iRFP showed higher fluorescence quantum yield than BV-bound iRFP. To facilitate the simple use of iRFP in fission yeast, we developed a plasmid for iRFP tagging of endogenous proteins at the C-terminus, novel genome integration vectors, and all-in-one plasmids carrying genes required for both the SynPCB system and iRFP-fused marker proteins.

## RESULTS

### iRFP does not fluoresce in fission yeast *Schizosaccharomyces pombe*

During the process of experiments, we accidentally found that iRFP did not fluoresce at all in fission yeast. We first tested whether iRFP was applicable to near-infrared imaging in fission yeast. We established a cell strain stably expressing nuclear localization signal (NLS)-iRFP-NLS (Miura et al. 2018) under the constitutive promoter *Padh1*. Two NLSs are fused with iRFP because the addition of a single NLS does not sufficiently localize the protein at the nucleus. No iRFP fluorescence was observed at an excitation wavelength of 640 nm (Fig. 1B). Because the iRFP requires BV as a chromophore for emitting fluorescence (Fig. 1A), we hypothesized that fission yeast could not metabolize BV intracellularly. Upon the addition of external BV, the nuclear iRFP fluorescence signal was recovered (Fig. 1B). The titration of BV concentration yielded a dose-dependent increase in iRFP fluorescence up to 125 μM (Fig. 1C). We next examined the kinetics of BV incorporation into fission yeast cells. Treatment with a high dose of BV (500 μM) gradually increased iRFP fluorescence until 24 h, suggesting slow uptake of BV in fission yeast cells (Fig. 1D). Since BV is produced from heme through HO, we searched for *HO* in the genomes of fission yeast and representative fungal species. As expected, we could not find any *HO* or *HO*-like gene in fission yeast (Fig. S1). Interestingly, *HO* and/or *HO*-like genes, which have been found from bacteria to higher eukaryotes, are frequently and sporadically lost in the representative fungal species (Fig. S1). Indeed, while iRFP has been widely used in the budding yeast, *Saccharomyces cerevisiae*, which retains an *HO* gene (Wosika et al. 2016; Yang Li et al. 2017; Geller et al. 2019; Tojima et al. 2019), there have been no studies using iRFP in the fission yeast, *S*.*pombe*. Taken together, these facts led us to conclude that iRFP does not fluoresce in fission yeast due to the lack of BV and *HO*.

### Development of novel stable knock-in plasmids: pSKI

The above results showed that the external supply of BV required a high dose and long-term incubation to realize iRFP fluorescence in fission yeast, which prompted us to seek an alternative route to iRFP fluorescence by introducing genes for the biosynthesis of BV. Before starting to develop the reconstitution system, we developed novel stable integration vectors that met our specific requirements—stable one copy integration into the genome, no effect on the auxotrophy of integrated cells, and distant integration loci for crossing strains—rather than using one of the previously developed integration systems (Keeney and Boeke 1994; Matsuyama et al. 2004; Maundrell 1993; Siam, Dolan, and Forsburg 2004; Fennessy et al. 2014; Kakui et al. 2015; Vještica et al. 2020). At first, we chose three gene-free loci on each chromosome at chromosome I positions 1,508,522 to 1,508,641 (near *mug165*, 1L), chromosome II positions 447,732 to 447,827 (near *pho4*, 2L), and chromosome III positions 1,822,244 to 1,822,343 (near *nup60*, 3R) (Fig. S2A). Next, we designed and developed plasmids that contain genes required for replication and amplification in *E. coli* (*Amp*, ori), the constitutive promoter *Padh1* or inducible promoter *Pnmt1*, a multiple cloning site (MCS), an *adh1* terminator, a selection marker cassette encoding an antibiotic-resistance gene for fission yeast, and homology arms connected with the one-cut restriction enzyme recognition site for plasmid linearization (Fig. S2B). Expected genomic integration with these vectors was confirmed by genomic PCR using primers designed to span the integration boundary (Fig. S2C). None of these integrations affected the bulk growth of fission yeast (Fig. S2D), and the protein expression levels from these three loci were comparable or moderately higher than that from the Z-locus (Fig. S2E). We named this series of plasmids using the prefix pSKI (<u>p</u>lasmid for <u>S</u>table <u>K</u>nock-<u>I</u>n, also see Table S1) and used them for the following experiments.

### PCB brightens iRFP more efficiently than BV in fission yeast

HO is the crucial enzyme in the BV biosynthesis pathway, catalyzing the linearization of tetrapyrrole (Fig. 2A). Therefore, we established fission yeast cells stably expressing HO1 and NLS-iRFP-NLS with pSKI and quantified the resulting iRFP fluorescence. As expected, the expression of HO1 derived from *Thermosynechococcus elongatus* BP-1 in mitochondria, where heme is abundant, demonstrated iRFP fluorescence, and the iRFP fluorescence was brighter than that achieved by the external addition of BV (Fig. 2B, second and third columns). Because HO1 is known to catalyze heme in the presence of reduced Fd (Rhie and Beale 1992), we next examined whether co-expression of HO1 and tFnr-Fd, a chimeric protein of truncated Fnr and Fd (Uda et al. 2020), would improve HO1-mediated iRFP fluorescence. However, the co-expression of HO1 and tFnr-Fd in mitochondria did not further enhance iRFP fluorescence as compared to the expression of only HO1 (Fig. 2B, sixth column), suggesting that authentic ferredoxin in fission yeast sufficiently supports the catalytic reaction through HO1.

**Fig 2.**
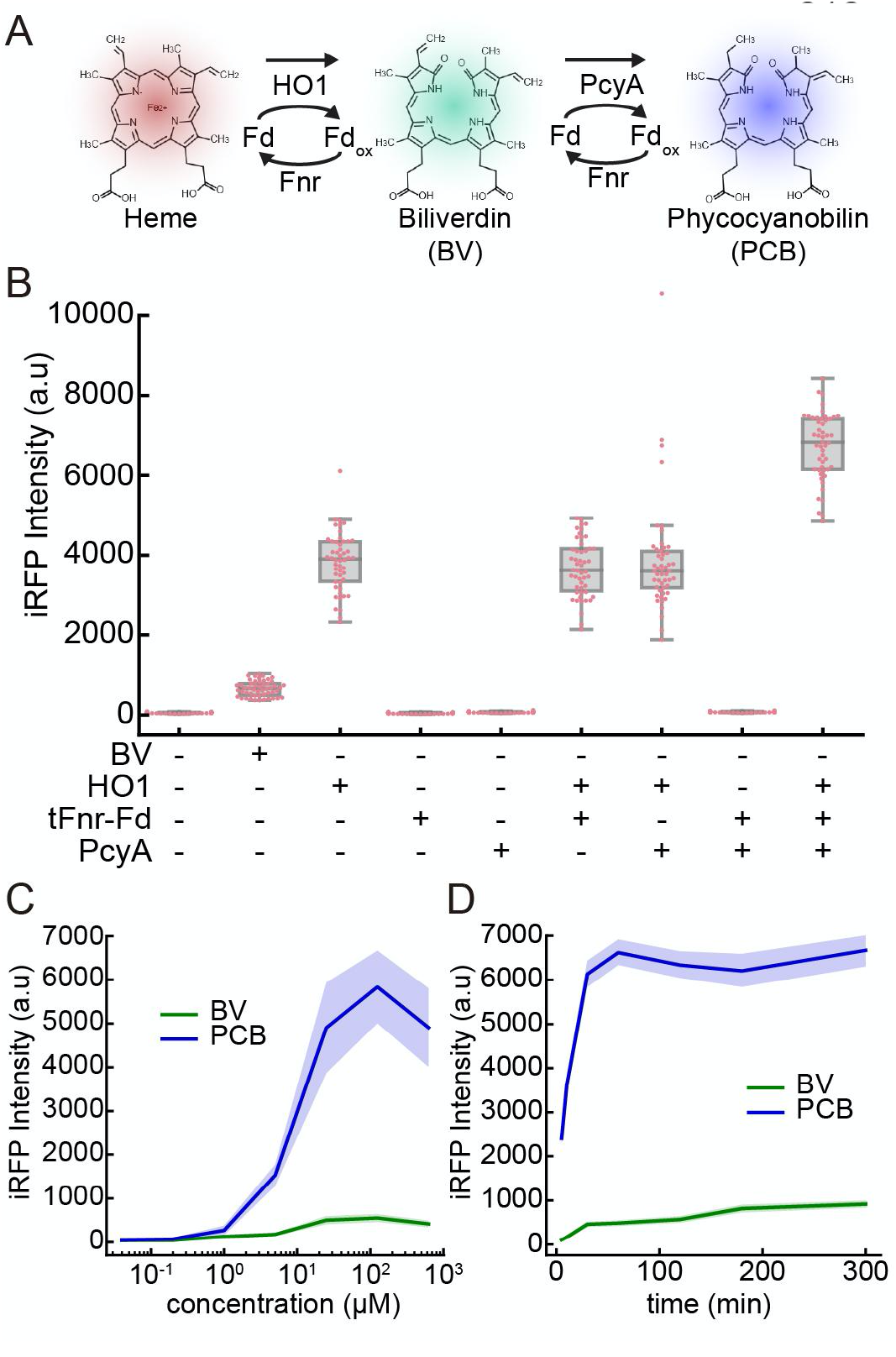
PCB brightens iRFP more efficiently than BV in fission yeast. (A) Schematic illustration of the PCB biosynthesis pathway. (B) Quantification of iRFP fluorescence in fission yeast cells expressing HO1, tFnr-Fd, and PcyA. Under the BV condition, cells were treated with 125 μM BV for 1 h at room temperature. Each dot represents iRFP fluorescence from a single cell with a boxplot, in which the box shows the quartiles of data with the whiskers denoting the minimum and maximum except for the outliers detected by 1.5 times the interquartile range (n = 50 cells). (C) Dose-response curve of iRFP fluorescence as a function of the levels of BV or PCB incorporation into fission yeast cells. Fission yeast cells were cultured in liquid YEA and incubated at room temperature for 3 h with the indicated concentration of BV or PCB (8 nM, 40 nM, 200 nM, 1 μM, 5 μM, 25 μM, 125 μM, and 625 μM). The lines and shaded areas indicate the averaged intensities and S.D., respectively (n = 50 cells). The decrease in iRFP intensity under 625 μM PCB or BV could be due to cell death and/or toxicity by the excess DMSO. (D) Time-course of iRFP fluorescence in response to BV or PCB treatment. Fission yeast cells were cultured in liquid YEA and treated with 125 μM BV or PCB at time zero. The lines and shaded areas indicate the averaged intensities and S.D., respectively (n = 50 cells).

Unexpectedly, in a series of experiments, we found a further increment in iRFP fluorescence by PCB (Fig. 2B, ninth column). When PcyA, the enzyme responsible for the production of PCB from BV, was co-expressed with HO1 and tFnr-Fd, the level of iRFP fluorescence was higher than other conditions (Fig. 2B, ninth column). To validate these results, we treated the cells expressing NLS-iRFP-NLS with purified PCB instead of BV. The addition of external PCB substantially outperformed the addition of BV with respect to iRFP fluorescence intensity (Fig. 2C and 2D). While the fluorescence intensities were quite different between PCB-bound iRFP (iRFP-PCB) and BV-bound iRFP (iRFP-BV), the effective concentration of the dose-response curve (Fig. 1C and 2C) and the kinetics of chromophore incorporation (Fig. 1D and 2D) were comparable between them.

### PCB yields brighter fluorescence as an iRFP chromophore than BV

The above data indicated the possibility that PCB might be a more suitable chromophore for iRFP than BV. To prove this hypothesis, we first examined whether the efficiency of holo-iRFP formation accounted for the difference in iRFP fluorescence between BV- and PCB-treated cells. PCB was added to the cells with HO1 expression, which exhibited constant intracellular production of BV. Therefore, iRFP has already formed a holo-complex with BV before attaching to PCB (Fig. 3A). Given that iRFP-PCB is brighter than iRFP-BV, we reasoned that HO1 expression attenuated the increase in iRFP fluorescence when the cells were further treated with purified PCB due to the competition between the PCB and already existing BV for binding to iRFP. As we expected, the addition of purified PCB hardly increased iRFP fluorescence in cells that had been expressing HO1, in spite of the dose-dependent increase in iRFP fluorescence by PCB treatment in cells not expressing HO1 (Fig. 3B and 3C). These observations reveal that almost all iRFP forms a holo-complex with BV when HO1 is expressed.

**Fig 3.**
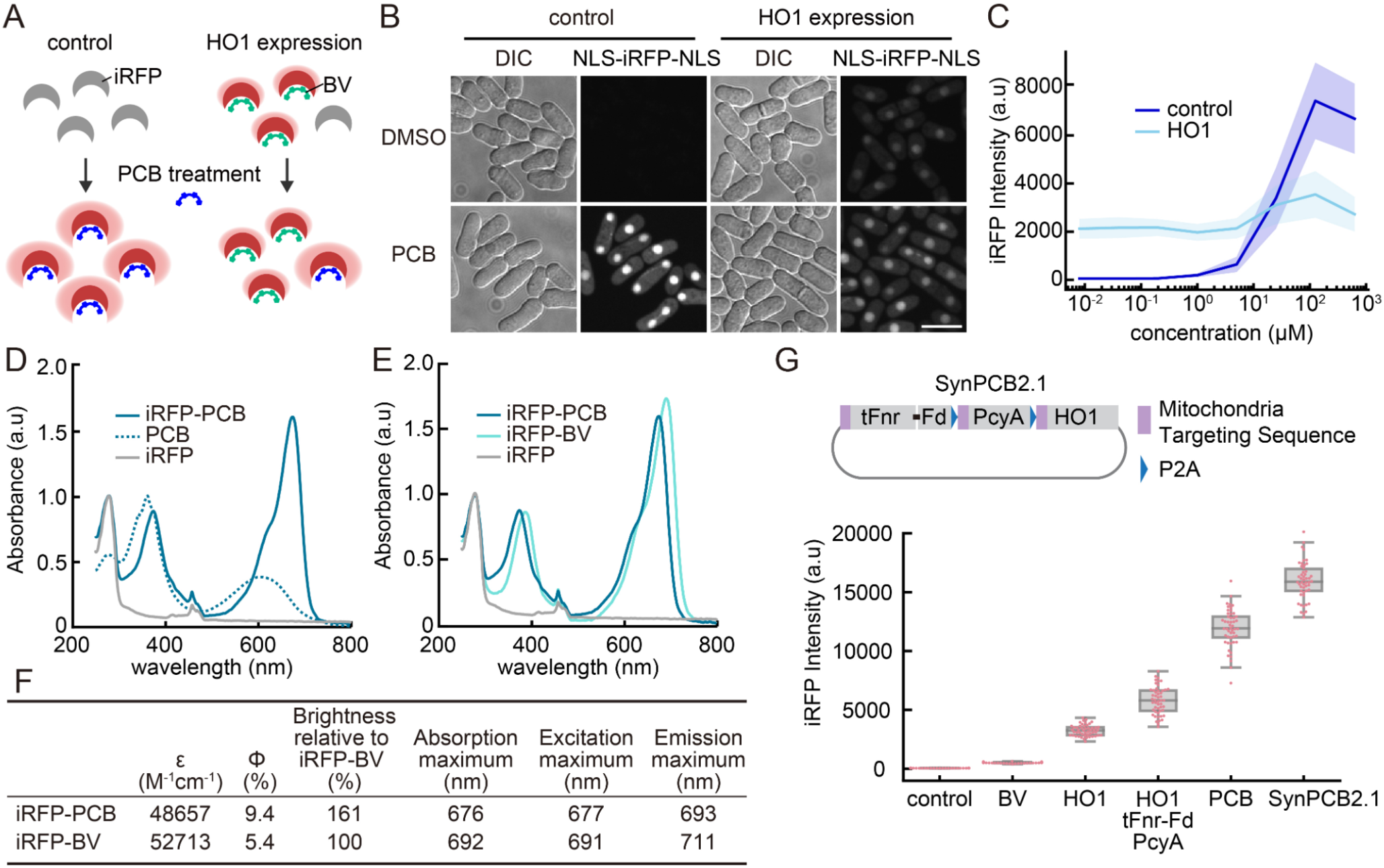
PCB yields brighter fluorescence as an iRFP chromophore than BV. (A) Schematic illustration of the experimental procedure. In control fission yeast cells, iRFP shows fluorescence upon the addition of PCB. In HO1 expressing cells, BV binds to iRFP as a chromophore before the addition of PCB. Therefore, BV competes with PCB for binding to iRFP. (B) Representative images of fission yeast expressing NLS-iRFP-NLS with or without external PCB (125 μM) treatment. Scale bar, 10 μm. (C) Dose-response curve of iRFP fluorescence as a function of the PCB concentration in a culture of fission yeast cells. Fission yeast cells were cultured in liquid YEA and incubated at room temperature for 1 h with the indicated concentration of PCB (8 nM, 40 nM, 200 nM, 1 μM, 5 μM, 25 μM, 125 μM, and 625 μM). The lines and shaded areas indicate the averaged intensities and S.D., respectively (n = 50 cells). (D) Normalized absorption spectra of PCB-bound iRFP (iRFP-PCB), free PCB, or iRFP. First, the spectra of iRFP-PCB and iRFP were normalized based on the absorbance at 280 nm (absorbance of protein), followed by normalization of the PCB spectrum by the absorbance at 375 nm. Of note, there is a spectrometer artifact at around 450 nm in all spectra. (E) Normalized absorption spectra of iRFP-PCB, BV-bound iRFP (iRFP-BV), and iRFP. The absorption spectra were normalized by the absorbance at 280 nm of each spectrum. Of note, there is a spectrometer artifact at around 450 nm in all spectra. (F) Summary of the fluorescence properties of iRFP-PCB and iRFP-BV *in vitro*. Φ and ε represent the fluorescence quantum yield and molar extinction coefficient, respectively. (G) (upper) Structure of the SynPCB2.1 plasmid expressing tFnr-Fd, PcyA, and HO1. These proteins are tagged with the mitochondria targeting sequence (MTS) at their N-termini and flanked by P2A, a self-cleaving peptide. (lower) Quantification of iRFP fluorescence under the indicated conditions. Cells were treated with 125 μM BV or PCB for 1 h at room temperature (second and fifth columns). Each dot represents iRFP fluorescence of a single-cell with a boxplot, in which the box shows the quartiles of data with the whiskers denoting the minimum and maximum except for the outliers detected by 1.5 times the interquartile range (n = 50 cells).

To understand why iRFP-PCB was brighter than iRFP-BV, we prepared recombinant iRFP expressed in *E. coli* and purified apo-iRFP (Filonov et al. 2011) (Fig. S3A). Apo-iRFP was mixed with PCB and BV to form holo-iRFP, *i*.*e*., iRFP-PCB and iRFP-BV, respectively (Fig. S3B). Binding of PCB to iRFP resulted in a change in the absorption spectrum from the free PCB (Fig. 3D). The absorbance maximum of iRFP-PCB was 10 nm blue-shifted from that of iRFP-BV (Fig. 3E). Fluorescence excitation and emission spectra were also 10 nm blue-shifted in iRFP-PCB compared to iRFP-BV (Fig. S3C and S3D). Notably, the fluorescence quantum yield of iRFP-PCB was nearly twice as high as that of iRFP-BV (0.094 vs. 0.054), while their molecular extinction coefficient values were comparable (Fig. 3F). These results were consistent with the previous work (Stepanenko et al. 2019). Based on these results, we concluded that iRFP forms a complex with PCB as a holo-form and that iRFP-PCB is brighter than iRFP-BV at the molecular level.

### SynPCB2.1 is ideal for iRFP imaging in fission yeast

For easy iRFP imaging using PCB as a chromophore, we introduced a system for efficient PCB biosynthesis, SynPCB2.1, in which the *tFnr-Fd, PcyA*, and *HO1* genes are tandemly fused with the cDNAs of the mitochondrial targeting sequences (MTS) at their N-termini, and flanked by self-cleaving P2A peptide cDNAs for multicistronic gene expression (Uda et al. 2020) (Fig. 3G). The single-cassette of SynPCB2.1 genes was knocked-in into cells expressing NLS-iRFP-NLS with a pSKI vector system, and expressed under the *adh1* promoter. The cells expressing SynPCB2.1 showed higher iRFP fluorescence than either cells treated with PCB or cells expressing the three genes individually (Fig. 3G). The protein expression levels of iRFP were comparable between the cells treated with BV and PCB, and cells expressing HO1 or SynPCB2.1 (Fig. S4). These results indicate that the chromophore formation of iRFP has little impact on the protein stability of iRFP in fission yeast.

To determine to what extent iRFP formed a complex with PCB or BV in cells, we quantified the fraction of fluorescent iRFP molecules by fluorescence correlation spectroscopy (FCS). FCS is the technique that exploits temporal fluctuation of fluorescent molecules in the confocal volume (1 fL), enabling to estimate the number of fluorescent molecules in the confocal volume and the diffusion coefficient (Shi et al. 2009; Sudhaharan et al. 2009; Kinjo, Sakata, and Mikuni 2011). For this purpose, fission yeast cells expressing iRFP fused with mNeonGreen (iRFP-mNeonGreen) were treated with BV or PCB or co-expressed with HO1 or SynPCB2.1, and subjected to FCS measurement to quantify the number of fluorescent iRFP and mNeonGreen molecules (Fig. S5A). The more iRFP forms a complex with the chromophore and fluoresces in cells, the more the ratio of the number of fluorescent iRFP molecules to the number of mNeonGreen measured by FCS approaches 1 (Fig. S5A). The cells expressing iRFP-mNeonGreen and SynPCB2.1, and the cells treated with PCB exhibited the ratio values of approximately 0.8 and 1.0, respectively (Fig. S5B and S5C), showing that 80-100% of iRFP forms a complex with PCB under these conditions. Importantly, HO1 expression resulted in the formation of a holo-iRFP complex with almost the same efficiency as SynPCB2.1 expression (Fig. S5C). Given the fact that the brightness of iRFP-BV was much weaker than that of iRFP-PCB (Fig. 3G), these results indicate that iRFP-PCB is a substantially brighter fluorescent protein than iRFP-BV in fission yeast. The external addition of BV resulted in lower values of iRFP to mNeonGreen ratio, suggesting that the BV incorporation is the rate-limiting step in fission yeast.

Next, we measured the emission spectrum of iRFP in a living cell to compare the fluorescence properties of iRFP-BV and iRFP-PCB. As for the emission spectrum *in vitro*, the cells showed a distinct emission spectrum between iRFP-PCB and iRFP-BV, namely, a blue-shifted emission spectrum of iRFP-PCB (Fig. S6A). A similar shift was observed when the emission spectrum of cells expressing SynPCB2.1 was compared to that of cells expressing HO1 (Fig. S6B and summarized in Fig. S6E). Importantly, cells separately expressing HO1, tFnr-Fd, and PcyA exhibited an intermediate emission spectrum, suggesting a mixture of iRFP-BV and iRFP-PCB in this cell line. The presence of iRFP-BV would explain why iRFP fluorescence by SynPCB2.1 was brighter than that generated by separate expression of the three enzymes in fission yeast (Fig. 3G). Moreover, the emission spectra obtained from living fission yeast cells demonstrated that iRFP-PCB was much brighter than iRFP-BV (Fig. S6C and S6D).

To explore the generality of the application of PCB and SynPCB2.1 system to other near-infrared fluorescent proteins, we measured fluorescence intensities of miRFP670 and miRFP703, which are derived from a different branch of bacteriophytochrome RpBphP1 (Shcherbakova et al. 2016), in fission yeast treated with BV or PCB or expressing SynPCB2.1 (Fig. S7). The fluorescence intensities of both miRFP670 and miRFP703 were enhanced by the addition of PCB and the expression of SynPCB2.1 compared to the addition of BV in a similar manner iRFP (Fig. S7). From these data, we concluded that PCB biosynthesis by SynPCB2.1 is suitable for imaging with near-infrared fluorescent proteins in fission yeast.

During iRFP imaging experiments, we found that PCB synthesized in fission yeast cells expressing SynPCB2.1 is leaked out of the cells and incorporated into the surrounding cells. To clearly show the PCB leakage, we co-cultured cells expressing only SynPCB2.1 and cells expressing only NLS-iRFP-NLS. While neither strains exhibited any fluorescence when cultured singly, NLS-iRFP-NLS emanated fluorescence when cells were co-cultured with the cells expressing SynPCB2.1 (Fig. S8B and S8C). The data indicate that in fission yeast, PCB is leaked into the extracellular space.

### iRFP imaging in fission yeast: Development of endogenous tagging and all-in-one integration systems

To further exploit the advantages of iRFP imaging in fission yeast, we first established C-terminal tagging plasmids based on a commonly used PCR-based tagging system (Longtine et al. 1998). The plasmids included an *iRFP* cassette followed by one of four different selection markers (Fig. 4A). By using these plasmids, we verified endogenous *iRFP* tagging to several genes, including *cdc2* (CDK, nucleus), *rpb9* (PolII, chromatin), *rpa49* (PolI, nucleolus), *swi6* (heterochromatin), *pds5* (cohesin), *cut11* (nuclear envelope), *mal3* (microtubule plus-end), *sfi1* (spindle pole body, SPB), *cox4* (mitochondria), and *cnx1* (endoplasmic reticulum, ER) with the expression of SynPCB2.1. All tested proteins showed the expected subcellular localization in fission yeast (Fig. 4B), although the signal-to-noise ratios depended on the expression level of the endogenously tagged proteins.

**Fig 4.**
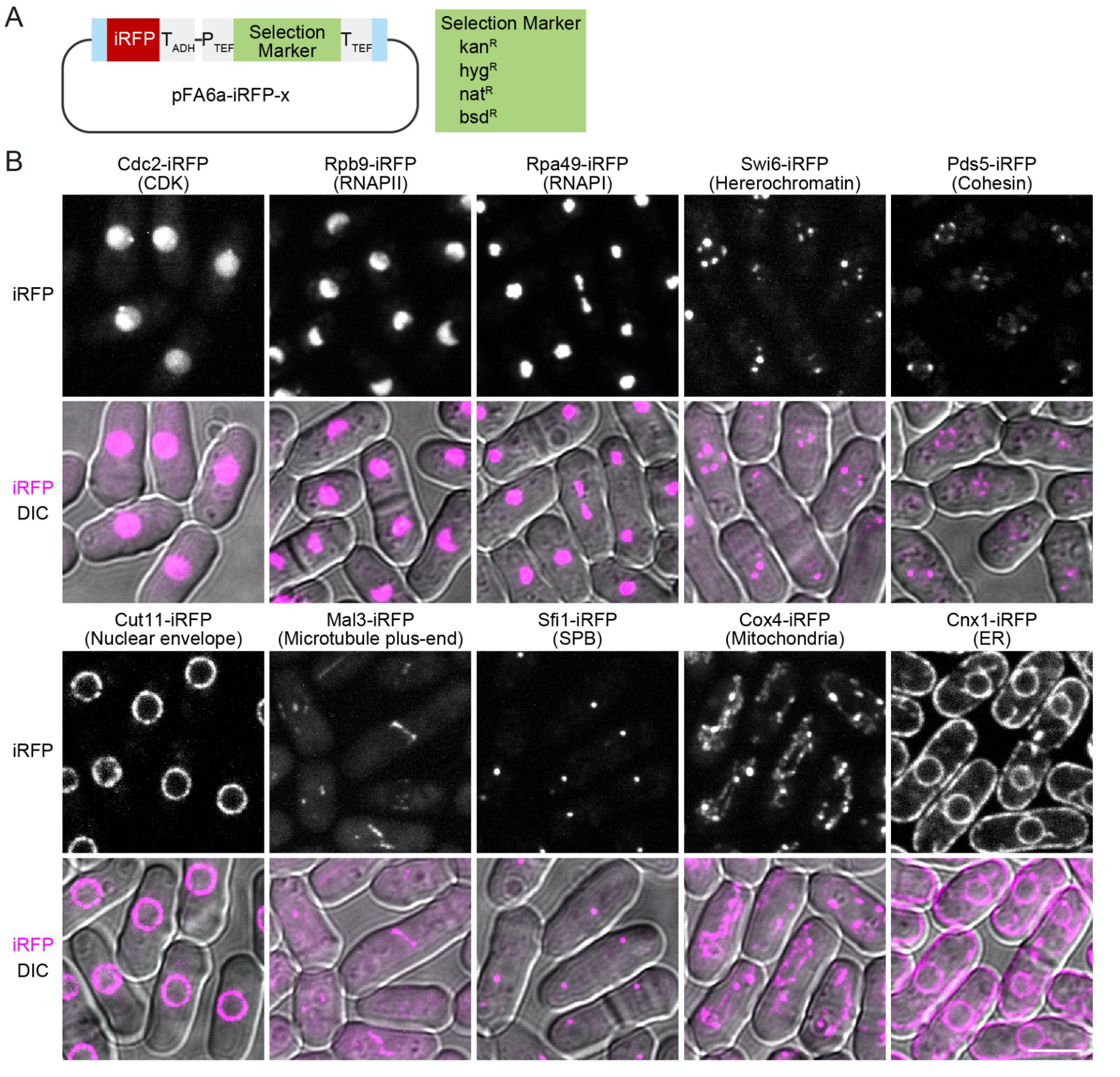
Visualization of endogenous proteins by iRFP in fission yeast. (A) Schematic illustration of the plasmid for iRFP tagging of endogenous proteins at the C-terminus. Cyan boxes indicate the common overlapping sequences (Longtine et al. 1998). The plasmid list is shown in Table S1. (B) The subcellular localization of endogenous proteins tagged with iRFP using pFa6a-iRFP. iRFP signals are shown in grayscale in the upper panels, and DIC images are merged with magenta iRFP signals and shown in the lower panels. Maximal projection images for iRFP are shown except for Cut11-iRFP and Cnx1-iRFP. Scale bar, 5 μm.

Second, we developed all-in-one plasmids carrying SynPCB2.1 and iRFP fusion protein genes to avoid a situation in which these two genes occupy two of the limited selection markers and integration loci. As a proof-of-concept, we introduced cDNA of Lifeact-iRFP (F-actin marker) or NLS-iRFP-NLS (nucleus marker) into the pSKI plasmid with the SynPCB2.1 gene cassette (Fig. 5A and 5B). Fission yeast transformed with these plasmids displayed the bright F-actin pattern, including actin patches, actin cables, and contractile ring (Fig. 5A) and nucleus (Fig. 5B). Taking full advantage of iRFP imaging with the SynPCB system in fission yeast, we established cells expressing five different proteins: The nucleus, plasma membrane, kinetochore, tubulin, and F-actin were labeled with NLS-mTagBFP2, Turquoise2-GL-ras1ΔN200, endogenous Mis12-mNeonGreen, mCherry-Atb2, and Lifeact-iRFP, respectively (Fig. 5C).

**Fig 5.**
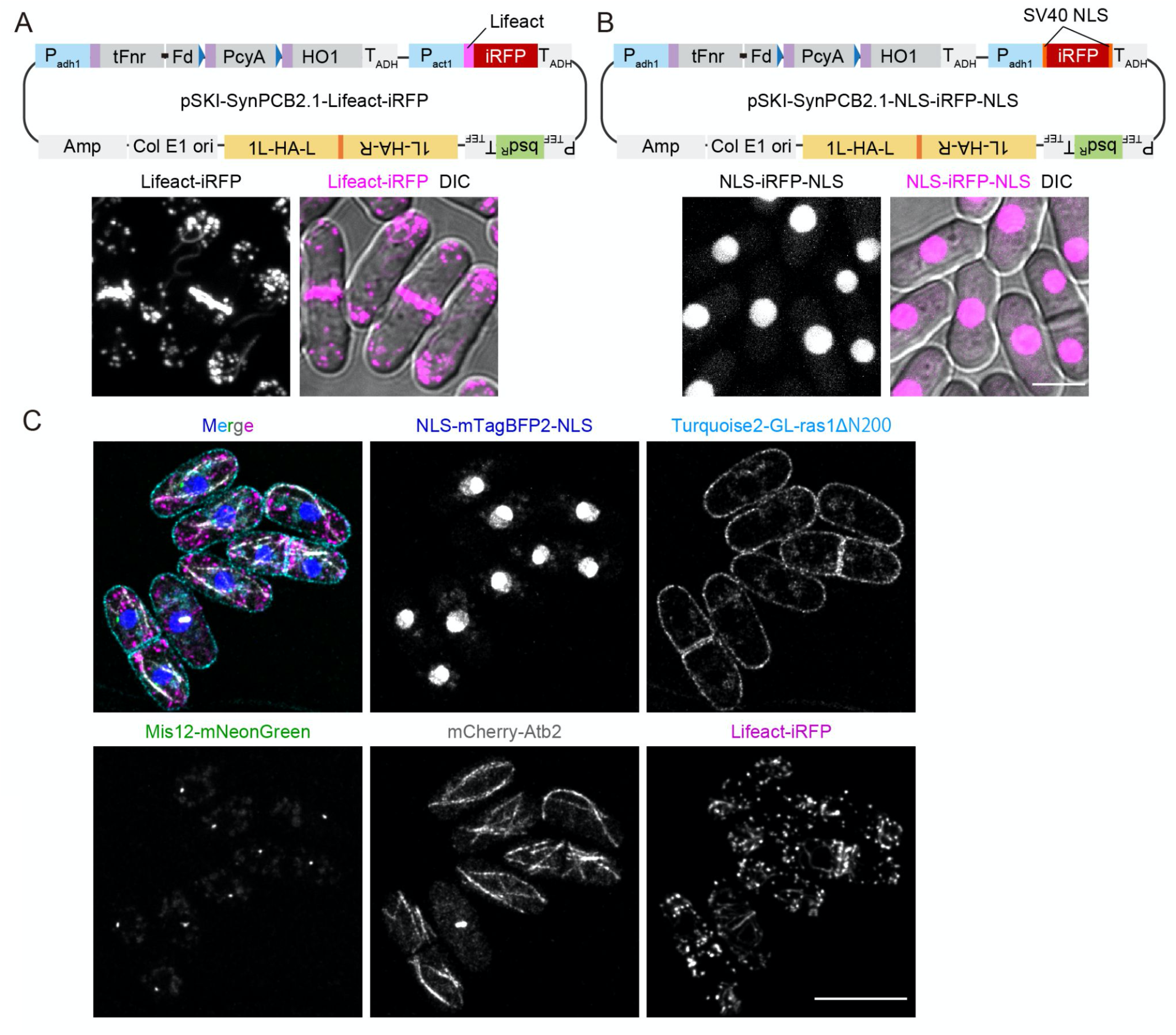
All-in-one plasmids for iRFP imaging. (A) (upper) Schematic illustration of 1L locus integration plasmids for the expression of SynPCB2.1 and Lifeact fused with iRFP (pSKI-SynPCB2.1-Lifeact-iRFP). (lower) Representative images of fission yeast expressing Lifact-iRFP are shown with the maximal intensity projection image and DIC- merged image. (B) (upper) Schematic illustration of 1L locus integration plasmids for the expression of SynPCB2.1 and NLS-iRFP-NLS (pSKI-SynPCB2.1-NLS-iRFP-NLS). (lower) Representative images of fission yeast expressing NLS-iRFP-NLS are shown with the maximal intensity projection image and DIC-merged image. Scale bar, 5 μm. (C) Multiplexed imaging of fission yeast expressing NLS- mTagBFP2-NLS (nucleus), Turquoise2-GL-ras1ΔN200 (plasma membrane), Mis12-mNeonGreen (kinetochore), mCherry-Atb2 (tubulin), and Lifeact-iRFP (F-actin). Maximal intensity projection images (except for Turquoise2-GL-ras1ΔN200; single z-section) and a merged image are shown. Scale bar, 10 μm.

### PCB can be used as a chromophore in mammalian cells

Finally, we tested whether PCB could be used as an iRFP chromophore in other organisms. HeLa cells expressing iRFP along with EGFP, an internal control for iRFP expression, were treated with external BV or PCB. PCB treatment increased the brightness of iRFP in HeLa cells to the same degree as BV treatment (Fig. S9A and S9B). BVRA KO HeLa cells displayed higher iRFP fluorescence than did parental HeLa cells, as reported previously (Kobachi et al. 2020), but did not show any change in iRFP fluorescence by BV or PCB treatment (Fig. S9B), probably because all iRFP molecules were occupied by BV. In contrast to fission yeast, the increment of iRFP fluorescence by PCB treatment was comparable to that by BV treatment in parental HeLa cells (Fig. S9B). Taken together, these results led us to conclude that PCB is applicable to iRFP imaging in mammalian cells, although it does not offer a significant advantage over BV.

## DISCUSSION

In this study, we demonstrated that iRFP does not fluoresce in fission yeast because of the lack of the BV-producing enzyme HO. Moreover, we found that PCB acts as a brighter chromophore for iRFP than BV both *in vitro* and in fission yeast expressing SynPCB2.1. Although PCB is not an authentic chromophore for iRFP nor the original RpBphP2, our data strongly suggested that PCB forms a fluorescent chromophore in iRFP. Finally, we developed endogenous iRFP tagging plasmids and all-in-one plasmids carrying SynPCB2.1 and iRFP marker proteins for the easy use of near-infrared imaging in fission yeast. As an alternative to external chromophore addition, the SynPCB2.1 system has potential advantages for iRFP imaging, including being fully genetically encoded and capable of providing even brighter iRFP fluorescence in fission yeast.

Our data indicate that PCB is more suitable as an iRFP chromophore than BV in fission yeast for several reasons. The first reason is that iRFP-PCB has a 2-fold higher fluorescence quantum yield than iRFP-BV *in vitro*. The second reason is that the excitation and emission spectra of iRFP-PCB are blue-shifted in comparison to those of iRFP-BV. This result is consistent with previous works describing the blue-shifted spectra of PCB (Rumyantsev et al. 2015; Loughlin et al. 2016). The blue-shifted spectra of iRFP-PCB possess favorable properties for most conventional confocal microscopes. Based on the emission and excitation spectrum (Fig. S3C), iRFP-PCB is approximately 1.3-fold more effectively excited by 640 nm of the excitation laser, and detected about 2.0-fold more efficiently with our emission filter (665-705 nm emission filter) in comparison to iRFP-BV. The third conceivable reason is the efficient chromophore formation. Indeed, RpBphP1-derived GAF-FP bound PCB 1.75-fold more efficiently than BV (Rumyantsev et al. 2015). Based on these data, the rough estimation yields 1.61*1.3*2*1.75 = 7.3-fold increase, which is comparable with the experimental results showing the 5∼10-fold increase in iRFP-PCB fluorescence compared to iRFP-BV (Figs. 2C, 2D, and 3G). In contrast to fission yeast, HeLa cells showed no difference in iRFP fluorescence between PCB and BV (Fig. S9). This could be partly due to the metabolism and culture conditions in mammalian cells, including synthesis of BV by endogenous HO1, degradation of BV and PCB by BVRA (Terry, Maines, and Lagarias 1993; Uda et al. 2017; Kobachi et al. 2020), and the presence of BV and bilirubin in the serum of the culture medium. Based on the results obtained by using fission yeast, we presume that the existence of BV within a HeLa cell and in the culture medium attenuates the increase in PCB-induced iRFP fluorescence. Moreover, other tetrapyrroles, such as primarily PPIX, could compete for iRFP with BV or PCB (Lehtivuori et al. 2013; Wagner et al. 2008).

The SynPCB system allows bright iRFP imaging without adding the external chromophores. This fact led us to consider that PCB might be applicable to other BV-based fluorescent proteins and optogenetic tools, as with miRFP670 and miRFP703, which exhibited increased fluorescence by SynPCB. Indeed, near-infrared fluorescent proteins that originate from allophycocyanin and cyanobacteriochrome, such as smURFP and iRFP670nano, respectively, exhibit high affinity to PCB because the original cyanobacteriochromes bind specifically to PCB (Rodriguez et al. 2016; Oliinyk et al. 2019). Bacteriophytochrome-based optogenetic tools using BV (Qian et al. 2020; Monakhov et al. 2020; Kaberniuk, Shemetov, and Verkhusha 2016; Redchuk et al. 2017) would be a potential target for the application of the SynPCB system. We should note that it is not clear whether PCB, instead of BV, increases the fluorescence brightness of these near-infrared fluorescent proteins and maintains the photoresponsive properties of these optogenetic tools. Fission yeast is an ideal model to assess phytochrome-based tools in a cell, such as the difference between BV and PCB as chromophores and the efficacy of genetically-encoded chromophore reconstruction because there is neither a synthetic nor a degradation pathway of BV in fission yeast.

We found that the *HO* homologue is frequently lost in fungal species, including the fission yeast during evolution (Fig. S1). In addition to fungi, *Caenorhabditis elegans*, one of the most popular model organisms, has shown very low, but not zero, BV-producing activity (Ding et al., 2017). Consistent with this fact, we could not find an *HO* homologue in the worm genome. The SynPCB system paves the way to utilizing iRFP for a broader range of organisms that lost an *HO* homologue during evolution. In addition, we recognized that PCB produced by SynPCB2.1 is leaked from the cells and taken up by surrounding cells, as evidenced by iRFP fluorescence (Fig. S8). It is possible that the same events take place under actual ecological conditions; some organisms may exploit tetrapyrroles produced by other organisms in order to render their own phytochromes functional. In fact, *Aspergillus nidulans* and *Neurospora crassa*, both of which lost an *HO* homologue in their genomes (Fig. S1), harbor phytochrome genes that are required for chromophores (Blumenstein et al. 2005; Froehlich et al. 2005). The exchanges of tetrapyrroles between living organisms might explain why the *HO* gene is sporadically lost in many organisms.

In this study, we have reported an iRFP imaging platform for fission yeast and a novel chromosome integration plasmid series, pSKI. The endogenous iRFP tagging system is based on a commonly used one, allowing anyone to introduce it quickly. The all-in-one plasmids carrying NLS-iRFP-NLS enable nuclear tracking without occupying green or red color fluorescence channels and automatic analysis of large-scale time-lapse images with nuclear translocation-type sensors (Regot et al. 2014). Further characterization and engineering will result in wide use of iRFP and phytochrome-based optogenetic tools in living organisms.

## MATERIALS AND METHODS

### Plasmids

The cDNAs of *PcyA, HO1, Fd*, and *Fnr* were originally derived from *Thermosynechococcus elongatus* BP-1 as previously described (Uda et al. 2020). The nucleotide sequence of these genes and SynPCB were optimized for human codon usage (see Benchling link; Table S1). The mitochondrial targeting sequence (MTS; MSVLTPLLLRGLTGSARRLP) was derived from human cytochrome C oxidase subunit VIII. The cDNAs were subcloned into vectors through conventional ligation with Ligation high Ver.2 (Toyobo, Osaka, Japan) or NEBuilder HiFi DNA Assembly (New England Biolabs, Ipswich, MA) according to the manufacturers’ instruction. The nucleotide sequence of mNeonGreen and Turquoise2-GL were optimized for fission yeast codon usage (see Benchling link; Table S1). The pSKI vectors include *Amp, colEI ori* (derived from pUC119), selection marker cassettes (derived from pFA6a-3FLAG-bsd, pFA6a-kan, pAV0587 (pHis5Stul-bleMX), pMNATZA1, and pHBCN1), *Padh1, Tadh1* (derived from pNATZA1), *Pnmt1, Tnmt1* (derived from pREP1), and MCSs (synthesized as oligo DNA (Fasmac)). To construct pSKI-SynPCB2.1-Lifeact-iRFP, *Pact1* (822 bp upstream of the start codon) was cloned from the fission yeast genome, and the cDNA of Lifeact was introduced by ligating annealed oligo DNAs. The cDNAs of miRFP670 and miRFP703 were obtained from pmiRFP670-N1 and pmiRFP703-N1 (gifts from Vladislav Verkusha, Addgene plasmids #79987 and #79988), and subcloned to obtain pMNATZA1-miRFP670 and pMNATZA1-miRFP703, respectively. pMNATZA1-iRFP-mNeonGreen was generated by inserting the cDNA of iRFP into the upstream of the mNeonGreen gene. All plasmids used in this study are listed in Table S1 with Benchling links, which include the sequences and plasmid maps.

### Reagents

Biliverdin hydrochloride was purchased from Sigma-Aldrich (30891-50MG), dissolved in DMSO (25 mM stock solution), and stored at –30°C. PCB was purchased from Santa Cruz Biotechnology (sc-396921), dissolved in DMSO (5 mM stock solution), and stored at –30°C. Of note, 625 μM PCB or 625 μM BV is insoluble in PBS solution and fission yeast culture medium, and 25 μM PCB or 25 μM BV is insoluble in mammalian cell culture medium because insoluble PCB or BV debris is observed.

### Fission yeast *Schizosaccharomyces pombe* strain and culture

All strains made and used in this study are listed in Table S2. The growth medium, sporulation medium, and other techniques for fission yeast were based on the protocol described previously (Moreno, Klar, and Nurse 1991) unless otherwise noted. The transformation protocol was modified from that of (Suga and Hatakeyama 2005). Genome integration by pSKI was confirmed by colony PCR with KOD One (TOYOBO) and the primers listed in Table S3. For the fluorescence microscope imaging, the fission yeast cells were concentrated by centrifugation at 3,000 rpm, mounted on a slide glass, and sealed by a cover glass (Matsunami).

### HeLa cell culture

HeLa cells were the kind gift of Michiyuki Matsuda (Kyoto University). BVRA KO HeLa cells have been established previously (Uda et al. 2017). HeLa cells were cultured in Dulbecco’s Modified Eagle’s Medium (DMEM) high glucose (Wako; Nacalai Tesque) supplemented with 10% fetal bovine serum (FBS) (Sigma-Aldrich) at 37°C in 5% CO_2_. For the live-cell imaging, HeLa cells were plated on CELLview cell culture dishes (glass bottom, 35 mm diameter, 4 components: The Greiner Bio-One). One day after seeding, transfection was performed with 293fectin transfection reagent (Thermo Fisher Scientific). Two days after transfection, cells were imaged with fluorescence microscopes. BV or PCB was added into the DMEM medium containing 10% FBS and cultured for 3 h at 37°C in 5% CO_2_.

### Measurement of the growth rate of fission yeast

Fission yeast cells were pre-cultured at 32 °C up to the optical density at 600 nm (OD600) of 1.0, followed by dilution to 1:20. A Compact Rocking Incubator Biophotorecorder TVS062CA (Advantec, Japan) was used for culture growth (32 °C, 70 rpm) and OD660 measurement. Growth curves were fitted by the logistic function (*x = K / (1 + (K/x*_*0*_ *-1)e*^*-rt*^)), and doubling time (*ln2/ r*) was calculated on Python 3 and Scipy.

### Protein purification

For the purification of His-tag fused iRFP, pCold-TEV-linker-iRFP was transformed into BL21(DE3) pLysS. *E. coli* (Promega, L1195) and selected on LB plates containing 0.1 mg/ml ampicillin at 37°C overnight. A single colony was picked up and inoculated into 2.5 ml liquid LB medium supplemented with 0.1 mg/ml ampicillin and 30 μg/ml chloramphenicol at 37°C overnight. The preculture was further inoculated into 250 mL liquid LB medium (1:100) containing ampicillin and chloramphenicol. The culture was shaken at 37°C for 2–4 h until the OD600 reached 0.6–1.0. The culture was then cooled to 18°C, and 0.25 mM Isopropyl β-D-1-thiogalactopyranoside (IPTG) (Wako, 094-05144) was added to induce the expression of His fused protein. After overnight incubation at 18°C, cells were collected and suspended into phosphate-buffered saline (PBS) (Takara, T900) containing 20 mM imidazole (Nacalai Tesque, 19004-22). Suspended cells were lysed sonication (VP-300N; TAITEC), followed by centrifugation to collect the supernatant. The supernatant was mixed with 250 μL Ni-NTA sepharose (Qiagen, 1018244), and incubated at 4°C for 2 h. Protein-bound beads were washed with PBS containing 20 mM imidazole, and proteins were eluted by the addition of 300 mM imidazole in PBS. Eluted fractions were checked by SDS-PAGE with a protein molecular weight marker, Precision Plus Protein™ All Blue Standards (Bio-Rad, #1610373), followed by CBB staining (BIOCRAFT, CBB-250) and detection by an Odyssey CLx system (Licor). Protein-containing fractions were dialyzed using a Slide-A-Lyzer Dialysis Cassette 3,500 MWCO (Thermo Scientific, 66110) to remove the imidazole. To concentrate the recombinant protein, Amicon ultra 3K 500 μL (Millipore, UFC500308) was used. To measure the protein concentration, Pierce™ BCA Protein Assay Kit (Thermo Scientific, 23227) was used. Purified His-iRFP in PBS solution was mixed with an excess amount (1:5 molar ratio) of BV or PCB dissolved in DMSO, followed by size exclusion chromatography with NAP-5 Columns (Cytiva, 17085301) to remove free BV or PCB.

### Characterization of *in vitro* fluorescence properties

The absorption of BV (100 μM), PCB (100 μM), and His-iRFP (12 μM) bound to chromophore was measured by a P330 nanophotometer (IMPLEN) with a 10 mm quartz glass cuvette (TOSOH, T-29M UV10). The absorption spectrum was measured in a wavelength range of 200 nm to 950 nm. For the measurements of absolute fluorescence quantum yield, BV or PCB bound His-iRFP (1 μM) in PBS was subjected to analysis with a Quantaurus-QY C11347-01 system (Hamamatsu Photonics). The excitation wavelength was 640 nm. For the measurements of excitation and emission spectra, BV- or PCB- bound His-iRFP (12 μM) was subjected to analysis with an F-4500 fluorescence spectrophotometer (Hitachi). The protein solution was excited in a wavelength range of 500 nm to 720 nm, and fluorescence at 730 nm was detected to measure the excitation spectrum. To measure the emission spectrum, the protein solution was excited at 640 nm, and fluorescence was detected in a wavelength range of 660 nm to 800 nm.

### Measurement of *in vivo* emission spectrum

The lambda-scan function of the Leica SP8 Falcon confocal microscope system was used for the measurement of the fluorescence emission spectrum. The excitation wavelength was fixed at 633 nm, and the 20 nm emission window was slid in 3 nm increments from 650 nm to 768 nm. Each emission spectrum was normalized by the peak emission value.

### Live-cell fluorescence imaging

Cells were imaged with an IX83 inverted microscope (Olympus) equipped with an sCMOS camera (ORCA-Fusion BT; Hamamatsu Photonics), an oil objective lens (UPLXAPO 100X, NA = 1.45, WD = 0.13 mm or UPLXAPO 60X, NA = 1.42, WD = 0.15 mm; Olympus), and a spinning disk confocal unit (CSU-W1; Yokogawa Electric Corporation). The excitation laser and fluorescence filter settings were as follows: excitation laser, 488 nm and 640 nm for mNeonGreen (or EGFP) and iRFP, respectively; excitation dichroic mirror, DM405/488/561/640; emission filters, 525/50 for mNeonGreen or EGFP, and 685/40 for iRFP (Yokogawa Electric). For the five colors multiplexed imaging, cells were imaged with Leica SP8 Falcon (Leica) equipped with an oil objective lens (HCPL APO CS2 100x/1.40 OIL). The excitation laser and fluorescence detectors settings were as follows: excitation laser, 405 nm, 470 nm, 488 nm, 560 nm, and 633 nm for mTagBFP2, Turquoise2-GL, mNeonGreen, mCherry, and iRFP, respectively; detector bandwidth, 420-450 nm, 480-500 nm, 500-550 nm, 580-650 nm, and 680-780 nm for mTagBFP2, Turquoise2-GL, mNeonGreen, mCherry, and iRFP, respectively. Images were obtained with 10 Z-slices of 0.5 μm intervals. Images were subjected to deconvolution by Lightning (Leica).

### Imaging analysis

All fluorescence imaging data were analyzed and quantified by Fiji (Image J). The background was subtracted by the rolling-ball method. Some images were obtained with 10–30 Z-slices of 0.2 μm intervals and shown as 2D images by the maximal intensity projection as noted in each figure legend. For the quantification of signal intensity, appropriate regions of interest (ROIs) were manually selected, and mean intensities in ROIs were measured.

### Fluorescence correlation spectroscopy (FCS) measurement and analysis

Time-series data of fluorescence fluctuation were obtained by Leica SP8 Falcon confocal microscope equipped with an objective lens HC PL APO 63x/1.20 W motCORR CS2, and were analyzed on Leica software essentially as described previously (Sadaie, Harada, and Matsuda 2014; Komatsubara et al. 2019). The structural parameter and effective confocal volume were calibrated using 500 nM Rhodamine 6G (TCI, cat #R0039) in DDW based on the result that the diffusion constant of Rhodamine 6G in DDW is 414 μm^2^/s at room temperature (C. B. Müller et al. 2008). The Rhodamine 6G solution was measured with 561 nm excitation and the emission from 580 nm to 700 nm. Structural parameter and the effective confocal volume were estimated as 3.70 and 0.616 fL, respectively. Fission yeast cells expressing iRFP-mNeonGreen fusion protein, whose molecule ratio of iRFP to mNeonGreen was 1:1, were measured as follow: excitation wavelength, 488 nm (mNeonGreen) and 640 nm (iRFP); emission window, from 500 nm to 620 nm (mNeonGreen) and from 680 nm to 768 nm (iRFP). Note that iRFP excitation laser power was increased when iRFP-BV was measured due to its dim fluorescence compared to iRFP-PCB. The time-series data of fluorescence fluctuation were obtained for 30 seconds, corrected by the photobleach correction algorithm on Leica FCS analysis software, and subject to the calculation of the auto-correlation function G(τ) on Leica FCS analysis software. The calculated auto-correlation functions were fitted with the equation concerning a single-component normal diffusion and triplet model on Leica FCS analysis software. The reciprocal of G (τ = 0), which is the amplitude of the correlation function, corresponds to the number of fluorescent molecules (N) in the confocal volume as *N* = 1/*G* (τ = 0). To estimate the fraction of holo-iRFP in all iRFPs, the number of fluorescent iRFP molecules was divided by that of fluorescent mNeonGreen molecules, assuming that all mNeonGreen formed chromophores.

### Analysis of *HO*-like sequences in representative species

We searched for *HO*-like sequences in representative fungal species using BLASTp (for details, see Table S4). We adopted human *HO1* (Uniprot P09601) and *S. cerevisiae HMX1* (Uniprot P32339) as the queries (e-value < 1e-5). The phylogenetic relationship is based on recent studies using multiple genes (Nguyen et al. 2017; Yuanning Li et al. 2021). We have manually drawn the evolutionary relationship among representative species (Nguyen et al. 2017) based on a recent genome-scale phylogeny (Yuanning Li et al. 2021), which is consistent with the current consensus view of the fungal tree of life (James et al. 2020). Since the results suggested sequence divergence among *HO1* homologues, we also used HO-like proteins of *Laccaria bicolor* and *Saitoella complicata* obtained from the BLASTp hits, although no additional sequence was found. Note that the absence in *Aspergillus nidulans* and the existence in *Candida albicans* are consistent with previous studies (Blumenstein et al. 2005; Pendrak et al. 2004). Concerning *C. elegans*, we searched for an *HO-*like sequence by the BLASTp interface provided on the WormBase website (http://www.wormbase.org, release WS280, date 20-Dec-2020, database version WS279). We used the same protein queries, *i*.*e*., human *HO1* and *S. cerevisiae HMX1*, although we obtained no hits (e-value < 1e-2).

## Acknowledgments

We thank all members of the Aoki Laboratory for their helpful discussions and assistance. The pCold-TEV plasmid was a kind gift of Dr. Koichi Kato (ExCELLS). We thank Dr. Takuya Norizuki for his critical reading and comments. Some fission yeast strains were provided by the National Bio-Resource Project (NBRP), Japan. We thank the Functional Genomics Facility of the NIBB Core Research Facilities for their technical support with fluorescence spectrometry.

## Competing Interests

The authors declare no competing or financial interest.

## Author contributions

Conceptualization: Y.G.; Data curation: Y.G., K.S., Y.K.; Formal analysis: Y.G., K.S., Y.K.; Funding acquisition: Y.K., M.K., Y.G., K.A.; Investigation: Y.G., K.S., Y.K., H.F, M.K.; Methodology: Y.G., K.S., Y.K., H.F, M.K.; Project administration: Y.G., K.A.; Resources: Y.G., K.S., H.F, M.K.; Supervision: K.A., Y.G.; Visualization: Y.G., K.S., Y.K.; Validation: Y.G., K.S., Y.K.; Writing - original draft: Y.G., K.S., Y.K., K.A.; Writing - review & editing: Y.G., K.S., Y.K., K.A.

## Funding

K.A. was supported by a CREST, JST Grant (JPMJCR1654), JSPS KAKENHI Grants (nos. 18H02444 and 19H05798), and the ONO Medical Research Foundation. Y.G. was supported by a JSPS KAKENHI Grant (no.19K16050), a Jigami Yoshifumi Memorial Research Grant, and a Sumitomo Research grant. Y.K. was supported by JSPS KAKENHI Grants (nos. 19K16207 and 19H05675).

## Notes

### Competing Interest Statement

The authors have declared no competing interest.

### Summary of Updates

We have responded to most of the reviewers' comments from Review Commons and revised the manuscript with additional new data.

