## Supplementary_Information for "Near-infrared imaging in fission yeast by genetically encoded biosynthesis of phycocyanobilin"

**Title:**

**Running Title:**

**iRFP imaging in fission yeast**

**Authors:**

Keiichiro Sakai[1,2,3], Yohei Kondo[1,2,3], Hiroyoshi Fujioka[4], Mako Kamiya[5], Kazuhiro Aoki[1,2,3,*], and Yuhei Goto[1,2,3,6*]

**Affiliations:**

[1]Quantitative Biology Research Group, Exploratory Research Center on Life and Living Systems (ExCELLS), National Institutes of Natural Sciences, 5-1 Higashiyama, Myodaiji-cho, Okazaki, Aichi 444-8787, Japan.

[2]Division of Quantitative Biology, National Institute for Basic Biology, National Institutes of Natural Sciences, 5-1 Higashiyama, Myodaiji-cho, Okazaki, Aichi 444-8787, Japan.

[3]Department of Basic Biology, School of Life Science, SOKENDAI (The Graduate University for Advanced Studies), 5-1 Higashiyama, Myodaiji-cho, Okazaki, Aichi 4ｇ44-8787, Japan.

[4]Graduate School of Pharmaceutical Sciences, The University of Tokyo, 7-3-1 Hongo, Bunkyo-ku, Tokyo 113-0033, Japan.

[5]Graduate School of Medicine, The University of Tokyo, 7-3-1 Hongo, Bunkyo-ku, Tokyo 113-0033, Japan

[6]Lead contact

*Corresponding authors

**Contact information**:

### Supplementary Figures

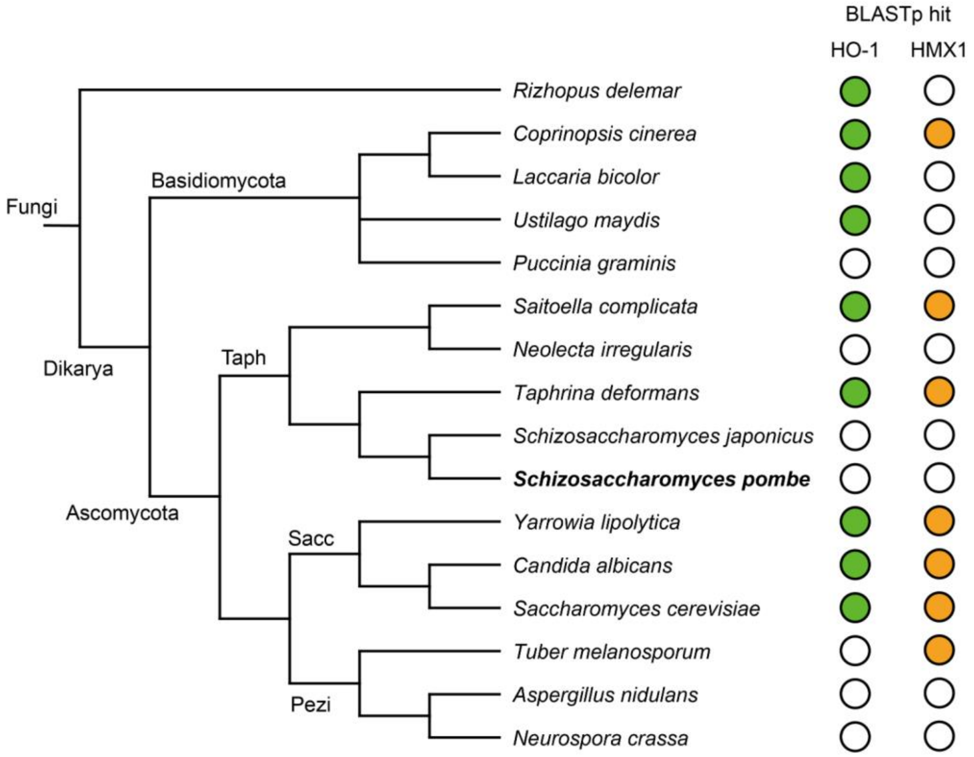


**Fig. S1. Phylogenetic distribution of HO-like sequences in fungal species**
Search results for heme oxygenase (HO)-like protein sequences in representative fungal species. Taph, Taphrinomycotina. Sacc, Saccharomycotina. Pezi, Pezizomycotina. The green circles indicate hits by BLASTp (e-value < 1e-5) with human HO1 as the query. The orange circles came from the same procedure except that *S.cerevisiae* HMX1 was the query.

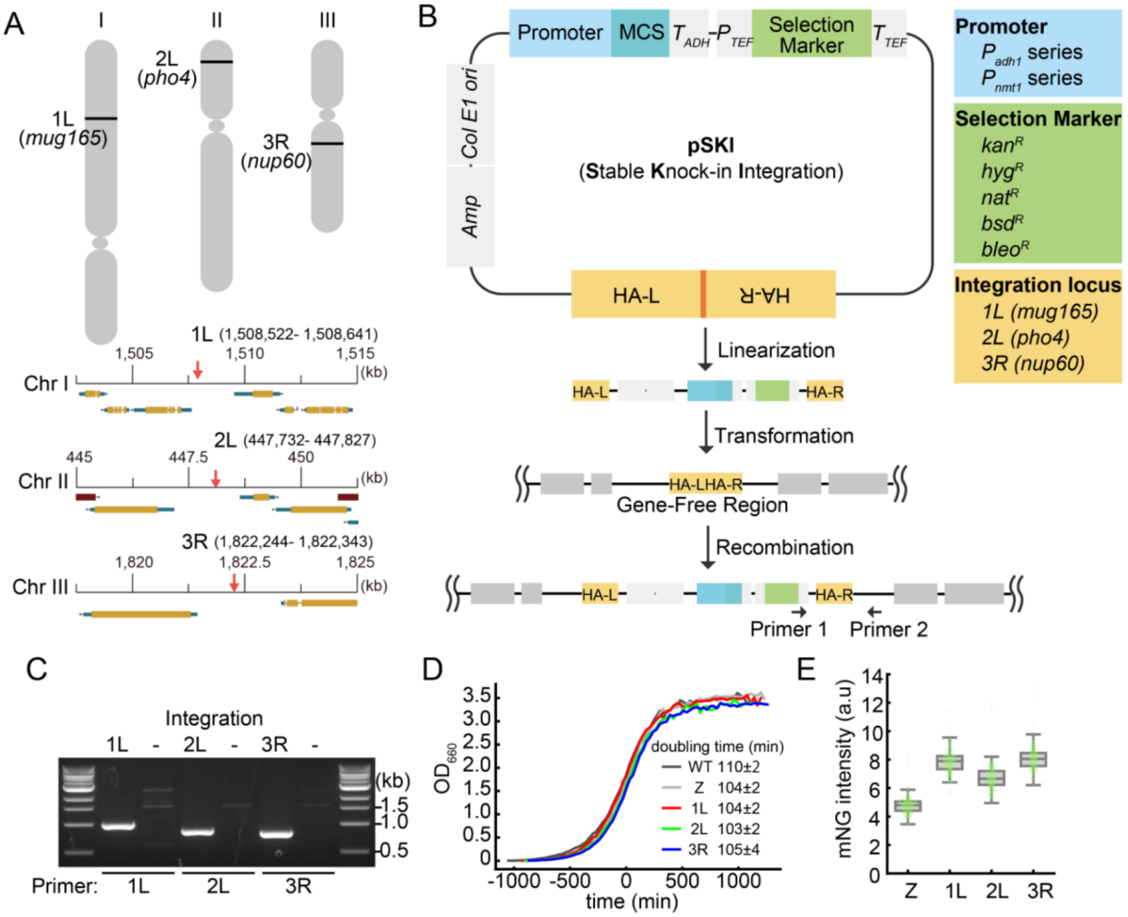


**Fig. S2. Novel chromosome integration plasmids, pSKI, for *Schizosaccharomyces pombe***
(A) Integration loci at the gene-free region of each chromosome (upper) and gene locations around integration loci (lower). Red arrows indicate the integration sites of each chromosome. (B) Plasmid map and integration procedure. The homology arm left and right (HA-L and HA-R) are connected with restriction enzyme recognition sites for linearization. Following linearization of the plasmid, DNA is introduced into the fission yeast cells by standard transformation procedures. Transformed DNA is integrated into the gene-free region through homologous recombination. The plasmid list is shown in Table S1. (C) Verification of the integration into each locus by PCR using primers 1 and 2 as indicated in panel B. Primer 1, which binds to the $T_{TEF}$ region, is common to all three loci (1L, 2L, and 3R), while primer 2, which binds to the outside of HA-R, is specific to each locus. (D) Representative growth curve of strains integrated with empty vectors of each locus. Mean doubling times are shown with the S.D. (n = 3, independent experiments). The calculation of the doubling time is described in the Materials and Methods. (E) Comparison of protein expression levels among 1L, 2L, 3R, and Z loci. mNeonGreen (mNG) was expressed from the indicated integration loci, and fluorescence intensity was quantified. Each dot represents the mNG fluorescence of a single cell with a boxplot, in which the box shows the quartiles of data with the whiskers denoting the minimum and maximum except for the outliers detected by 1.5 times the interquartile range (n > 150 cells).

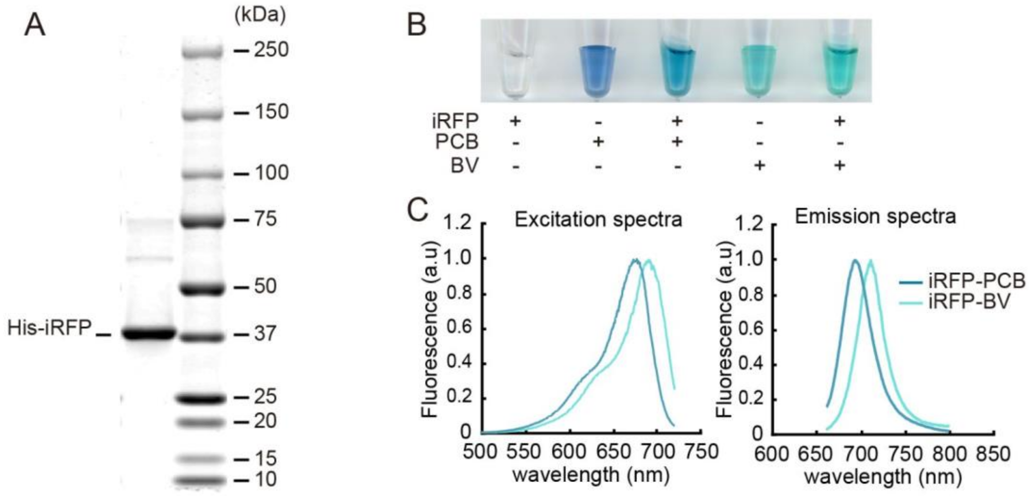


**Fig. S3. PCB binds to iRFP *in vitro*.**
(A) A representative image of CBB staining of purified recombinant His-iRFP (39 kDa). (B) A photograph of the purified His-iRFP, free PCB, PCB-bound His-iRFP, free BV, and BV-bound His-iRFP. Of note, unbound BV or PCB was eluted during the process of the size exclusion chromatography. (C) Excitation (left) and emission (right) spectra of His-iRFP bound to BV or PCB. The fluorescence intensities are normalized by the peak intensities.

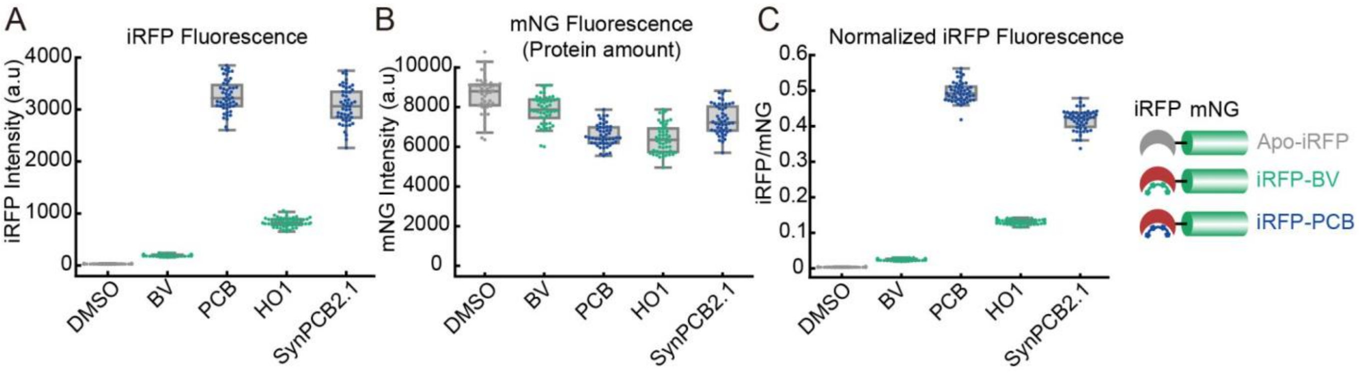


**Fig. S4. Quantification of expression levels of iRFP**

(A) iRFP fluorescence and (B) mNG fluorescence were quantified in fission yeast cells expressing iRFP-mNG. The cells were treated with DMSO, 125 μM BV, or 125 μM PCB for 2 h at room temperature (DMSO, BV, PCB) or co-expressed with HO1 or SynPCB2.1 (HO1, SynPCB2.1). iRFP fluorescence intensities were normalized by mNG fluorescence intensities (C). iRFP fluorescence (A), mNG fluorescence (B), and the ratio of iRFP fluorescence intensity to mNG fluorescence intensity (C) of each cell are plotted (Gray: Apo-iRFP, Green: iRFP-BV, Blue: iRFP-PCB) with a boxplot, in which the box shows the quartiles of data with the whiskers denoting the minimum and maximum except for the outliers detected by 1.5 times the interquartile range (n = 50 cells).

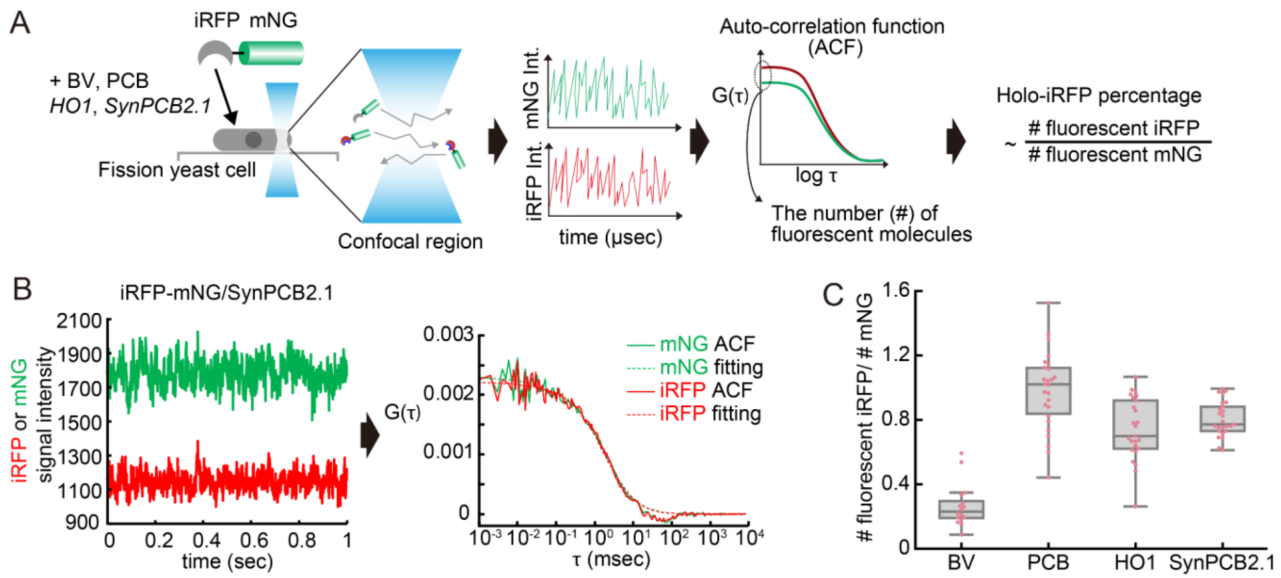


**Fig. S5. Quantification of the fraction of holo-iRFP by fluorescence correlation spectroscopy (FCS)**

(A) Schematic diagram of fluorescence correlation spectroscopy (FCS) analysis. Cells expressing iRFP-mNG were treated with 125 μM BV, or 125 μM PCB for 3 h at room temperature or co-expressed with SynPCB2.1, and subjected to FCS measurement. Temporal fluctuations of iRFP and mNG fluorescence in a tiny confocal volume were measured, and auto-correlation functions were calculated from the time-series data. The y-axis intercept of the auto-correlation functions ($G(\tau = 0)$) is inversely correlated to the number of fluorescence molecules in a confocal volume. Hence, the fraction of fluorescent holo-iRFP can be estimated by comparing the number of fluorescent iRFP with that of mNG molecules (see details in Materials and Methods). (B) Representative FCS data obtained from SynPCB2.1 expressing cells. The left graph shows the raw data of fluorescence fluctuation of iRFP (red) and mNG (green). The right graph represents the calculated auto-correlation functions (ACF, lines) and the fitted curves (fitting, dashed lines) of iRFP (green) and mNG (green) (see details in Materials and Methods). (C) The fraction of fluorescent holo-iRFP molecules in a living fission yeast cell was estimated by calculating the ratio of the number of fluorescent iRFP molecules to that of fluorescent mNG molecules. The ratio value of each cell under the indicated conditions is plotted with a boxplot, in which the box shows the quartiles of data with the whiskers denoting the minimum and maximum except for the outliers detected by 1.5 times the interquartile range ($n > 15$ cells).

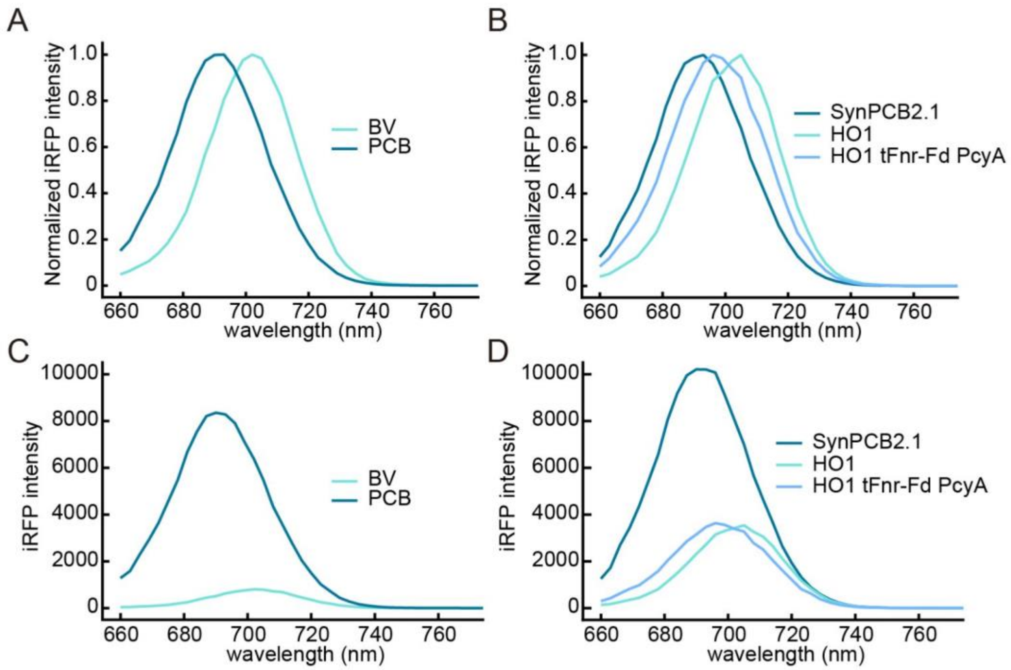


E

| | Emission maximum (nm) |
|---|---|
| BV | 702 |
| PCB | 690 |
| SynPCB2.1 | 690 |
| HO1 | 705 |
| HO1 tFnr-Fd PcyA | 696 |

**Fig. S6. Emission spectra of iRFP-BV or iRFP-PCB *in vivo***
(A) Normalized emission spectra of iRFP-BV and iRFP-PCB in fission yeast cells expressing NLS-iRFP-NLS treated with BV (125 μM) and PCB (125 μM), respectively. Fission yeast cells were excited by a 640 nm laser light source, and emission was measured every 3 nm with a 20 nm window. The fluorescence intensity at each window was averaged from over 10 cells. The fluorescence intensities were divided by the peak intensity for the normalization. (B) Normalized emission spectra of fission yeast cells expressing SynPCB2.1, HO1, or three genes (HO1, tFnr-Fd, and PcyA). (C and D) Raw emission spectra of panel A (C) and panel B (D). (E) Summary table of emission peaks *in vivo*.

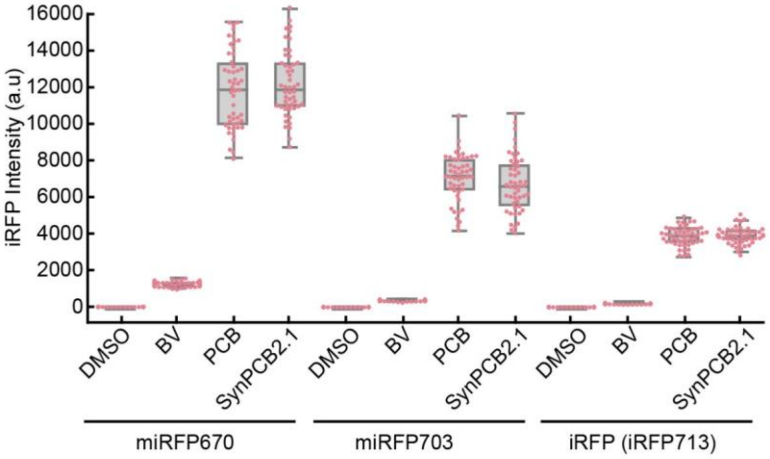


**Fig. S7. PCB is better chromophore to miRFPs than BV**

Quantification of miRFP fluorescence in fission yeast cells. Cells expressing miRFP670, miRFP703, or iRFP (= iRFP713) were treated with 125 µM BV or 125 µM PCB for 3 h at room temperature, or co-expressed with SynPCB2.1. Fluorescence intensity of miRFP670, miRFP703, or iRFP of each cell is plotted with a boxplot, in which the box shows the quartiles of data with the whiskers denoting the minimum and maximum except for the outliers detected by 1.5 times the interquartile range (n = 50 cells).

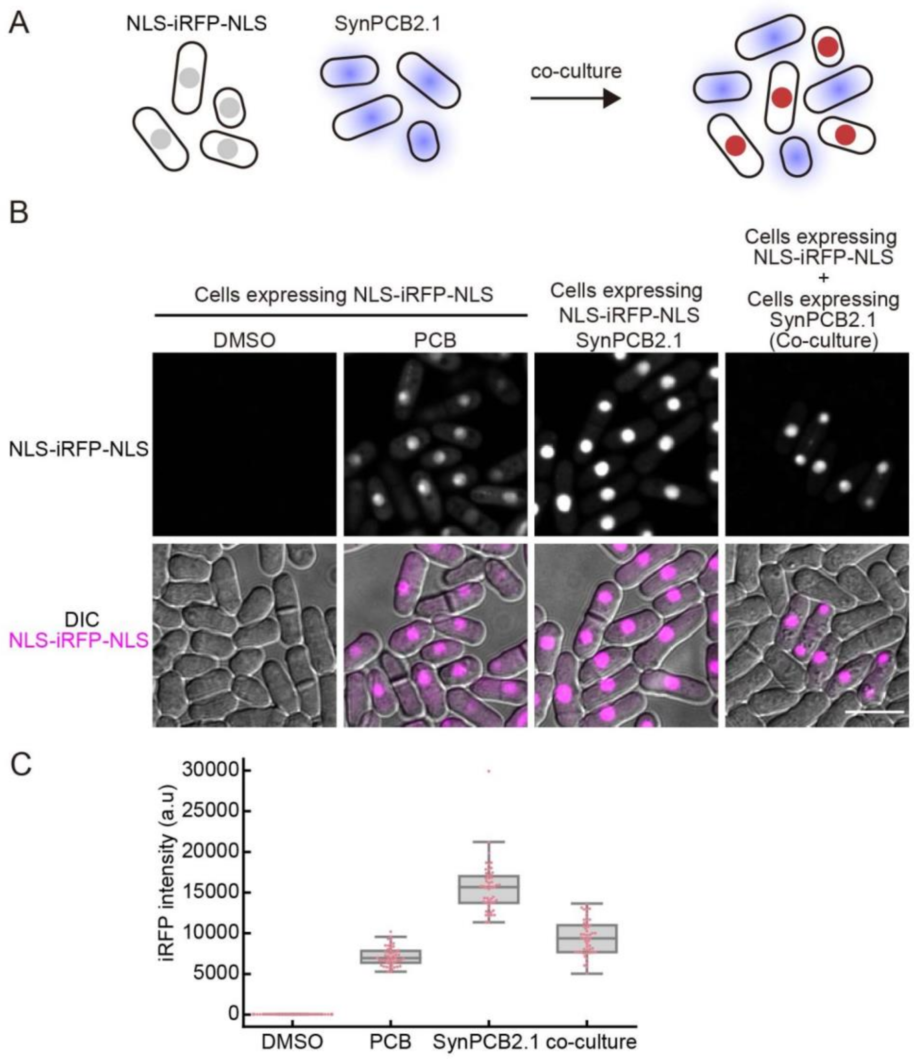


**Fig. S8. PCB is leaked from fission yeast cells expressing SynPCB2.1.**
(A) Schematic illustration of the co-culture experiment. Neither cells expressing only SynPCB2.1 nor cells expressing only NLS-iRFP-NLS exhibit iRFP fluorescence. NLS-iRFP-NLS exhibits iRFP fluorescence when the two cell lines are co-cultured due to the leaking of PCB from cells into the culture media. (B) Representative images of cells expressing NLS-iRFP-NLS treated with DMSO (first column), PCB (125 μM, second column), co-expression of SynPCB2.1 (third column), and co-culture with cells expressing SynPCB2.1 (fourth column). Scale bar, 10 μm. (C) Quantification of NLS-iRFP-NLS signal intensity in panel B. Single-cell fluorescence intensities is shown by dots with boxplots, in which the box shows the quartiles of data with the whiskers denoting the minimum and maximum except for the outliers detected by 1.5 times the interquartile range (n = 50 cells).

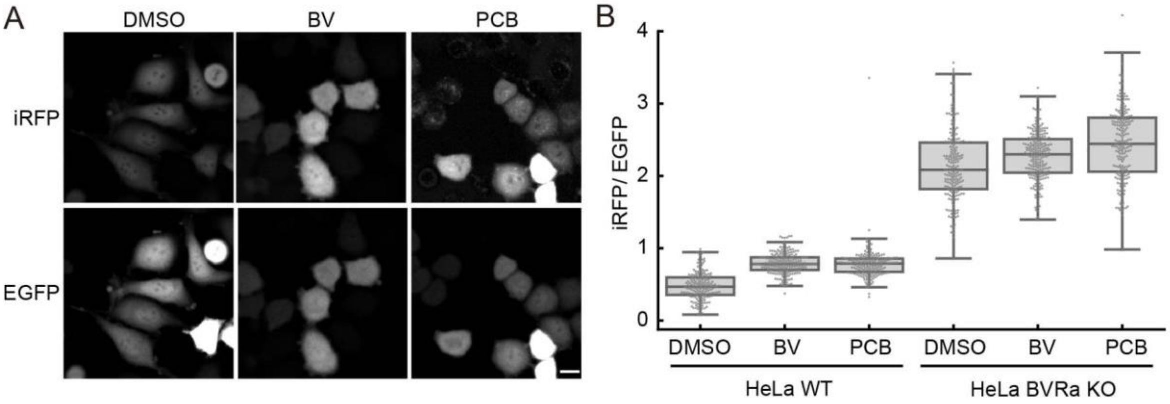


**Fig. S9. iRFP fluorescence in mammalian cells treated with PCB or BV**
(A) Representative images of parental HeLa cells expressing iRFP-P2A-EGFP treated with DMSO, BV (25 μM), or PCB (25 μM). Scale bar, 10 μm. Of note, 25 μM BV or PCB remained unsolved in the cultured medium for mammalian cells, and therefore chromophores in the medium could be saturated under these conditions. (B) Quantification of iRFP/EGFP values in HeLa cells and HeLa/BVRA KO cells under the indicated conditions. Each dot represents a data point from a single cell with a boxplot, in which the box shows the quartiles of data with the whiskers denoting the minimum and maximum except for the outliers detected by 1.5 times the interquartile range (n > 170 cells).

**Table S1. Plasmid list**

| Plasmid name | Description | Source | Benchling Link |
|---|---|---|---|
| pSKI-KAN-1L-A1-M | pSKI | this study | https://benchling.com/s/seq-G3P9JqxX51ObX5Sz96MA |
| pSKI-NAT-1L-A1-M | pSKI | this study | https://benchling.com/s/seq-8eBCjMRKikhCkkluJGhW |
| pSKI-BSD-1L-A1-M | pSKI | this study | https://benchling.com/s/seq-HhBJcOmn70PBjUtaRdFR |
| pSKI-BLE-1L-A1-M | pSKI | this study | https://benchling.com/s/seq-bhdt76l7rpG8s7St32bv |
| pSKI-KAN-2L-A1-M | pSKI | this study | https://benchling.com/s/seq-TreXp3COB6wHS3lOhASP |
| pSKI-NAT-2L-A1-M | pSKI | this study | https://benchling.com/s/seq-HLDrXgZxwWWQzFpCQmxk |
| pSKI-BSD-2L-A1-M | pSKI | this study | https://benchling.com/s/seq-eDFPR3eDDbwOpHn3wELo |
| pSKI-BLE-2L-A1-M | pSKI | this study | https://benchling.com/s/seq-tU4cqJD4RsS3y9AFcgvD |
| pSKI-KAN-3R-A1-M | pSKI | this study | https://benchling.com/s/seq-QLwzrxGXyx0TpSjNMzAH |
| pSKI-NAT-3R-A1-M | pSKI | this study | https://benchling.com/s/seq-Hvoe4y6Rfz9HRKKXmM8Q |
| pSKI-HYG-3R-A1-M | pSKI | this study | https://benchling.com/s/seq-EkO6jh5uMmuXF5brQFBq |
| pSKI-BSD-3R-A1-M | pSKI | this study | https://benchling.com/s/seq-NUrSW450XbywsQzxnyX9 |
| pSKI-BLE-3R-A1-M | pSKI | this study | https://benchling.com/s/seq-G4LydqOGtHxT2v7SHikY |
| pSKI-KAN-1L-N1 | pSKI | this study | https://benchling.com/s/seq-lajqCWRwNsuhe2ckb9nk |
| pSKI-NAT-1L-N1 | pSKI | this study | https://benchling.com/s/seq-X1w13cdTQVw4wgGpBfZR |
| pSKI-BSD-1L-N1 | pSKI | this study | https://benchling.com/s/seq-QQZRlvatv7zHfuA7bMnI |
| pSKI-BLE-1L-N1 | pSKI | this study | https://benchling.com/s/seq-MfQgtyoBCoIVSDIujzjg |
| pSKI-KAN-2L-N1 | pSKI | this study | https://benchling.com/s/seq-CFloX6NrLvuJrnI5JaPD |
| pSKI-NAT-2L-N1 | pSKI | this study | https://benchling.com/s/seq-YbDO5EX4oqDfJJguRrf0 |
| pSKI-HYG-2L-N1 | pSKI | this study | https://benchling.com/s/seq-8e48M8moxUXOqsybHbYO |
| pSKI-BSD-2L-N1 | pSKI | this study | https://benchling.com/s/seq-a5dfK1ki5ow43Px6o0tl |
| pSKI-BLE-2L-N1 | pSKI | this study | https://benchling.com/s/seq- |

| | | | |
|---|---|---|---|
| | | | KocnJoOdrJwaDPHHGeke |
| pSKI-KAN-3R-N1 | pSKI | this study | https://benchling.com/s/seq-BZmnME31WCynpEwW7GfH |
| pSKI-NAT-3R-N1 | pSKI | this study | https://benchling.com/s/seq-kVuyYx3nHglfwjYI0MYPP |
| pSKI-BSD-3R-N1 | pSKI | this study | https://benchling.com/s/seq-7M97ws9s2eY5ZfqXlQBq |
| pSKI-HYG-3R-N1 | pSKI | this study | https://benchling.com/s/seq-1eL7D3ki8KLvb3Uh3etM |
| pSKI-BLE-3R-N1 | pSKI | this study | https://benchling.com/s/seq-kMwFYBeWLuaYDoegFvdl |
| pFA6a-iRFP-kan | iRFP C-terminal tagging | this study | https://benchling.com/s/seq-bkYIYdFO0LlSWLRgqKf2 |
| pFA6a-iRFP-hyg | iRFP C-terminal tagging | this study | https://benchling.com/s/seq-emc65aNbjBFYkdt8Ooib |
| pFA6a-iRFP-nat | iRFP C-terminal tagging | this study | https://benchling.com/s/seq-hb5KLMGZXpgTFmQN9MH5 |
| pFA6a-iRFP-bsd | iRFP C-terminal tagging | this study | https://benchling.com/s/seq-MB0yvsIASTKHVyJnvflU |
| pFA6a-mNeonGreen (S.p codon optimized)-kan | mNG C-terminal tagging | this study | https://benchling.com/s/seq-r5vPvd2m4SeMm6VxVhiN |
| pSKI-KAN-1L-A1-M-SynPCB2.1 | SynPCB2.1 | this study | https://benchling.com/s/seq-iAht1qGQ6taledZRBjhy |
| pSKI-NAT-1L-A1-M-SynPCB2.2 | SynPCB2.1 | this study | https://benchling.com/s/seq-6Ds1klA3foLeFGMjNRY4 |
| pSKI-BSD-1L-A1-M-SynPCB2.1 | SynPCB2.1 | this study | https://benchling.com/s/seq-t5O2P2WzKmrmTYCcEtUk |
| pSKI-BLE-1L-A1-M-SynPCB2.1 | SynPCB2.1 | this study | https://benchling.com/s/seq-TeeZbKcTCjCZxRoCQ9ua |
| pSKI-BSD-1L-A1-M-SynPCB2.1-Lifeact-iRFP | pSKI-SynPCB2.1-Lifeact-iRFP | this study | https://benchling.com/s/seq-kazB0PhzppnUokAcnZBp |
| pSKI-BSD-1L-A1-M-SynPCB2.1-Tadh1-Padh1-NLS-iRFP-NLS-Tadh1 | pSKI-SynPCB2.1-NLS-iRFP-NLS | this study | https://benchling.com/s/seq-gGVpZDolap5t4mj7bGFK |
| pSKI-BSD-2L-A1-M-NLS-iRFP-NLS | | this study | https://benchling.com/s/seq-obPOLIX96PllBreCKNXY |
| pSKI-NAT-1L-A1-M-MTS-HO1 | | this study | https://benchling.com/s/seq-A38a1P9LQU1mhRfdWPBb |
| pSKI-NAT-3R-A1-M-MTS-btFnr-bFd | | this study | https://benchling.com/s/seq-6MjlWH76M7373w9WG02R |
| pMNATZA1-MTS-PcyA | | this study | https://benchling.com/s/seq-sRCmN6qpFCs6Dp5yVqzO |
| pSKI-KAN-1L-A1-M-MTS-HO1 | | this study | https://benchling.com/s/seq-U8xwbETwRXjlZmqHhQ2O |
| pSKI-KAN-3R-A1-M-MTS-btFnr-bFd | | this study | https://benchling.com/s/seq-SuqW1exP3iAJHvnDvxeD |

| | | | |
|---|---|---|---|
| pMNATZA1 | | this study | https://benchling.com/s/seq-F7zeXNTMu66Gs0zKvIpx |
| pMNATZA1-spmNeonGreen | | this study | https://benchling.com/s/seq-eNXfcf6oImjRKuJpX9k5 |
| pSKI-NAT-1L-A1-M-spmNeonGreen (S.p codon optimized) | | this study | https://benchling.com/s/seq-5RTHLQjGhwtHW5hpF9XC |
| pSKI-NAT-2L-A1-M-spmNeonGreen (S.p codon optimized) | | this study | https://benchling.com/s/seq-5KTJy6ykRRzXgNaRUUeV |
| pSKI-NAT-3R-A1-M-spmNeonGreen (S.p codon optimized) | | this study | https://benchling.com/s/seq-8eBCjMRKikhCkkluJGhW |
| pNATZA15-mCherry-atb2 | | this study | https://benchling.com/s/seq-0XMOLayGLNMT0Jx5xFa8 |
| pHBCA11-NLS-mTagBFP2-NLS | | this study | https://benchling.com/s/seq-vsO72fejJfzTcSFPqF5X |
| pMBLE-3R-A1-Turquoise2-GL-ras1delN200 | | this study | https://benchling.com/s/seq-KXyKpjnMwc3D1kstue68 |
| pCold-TEV-linekr-iRFP | | this study | https://benchling.com/s/seq-2HCLXjM6wnXr2mMIa5QH |
| pCAGGS-iRFP-P2A-EGFP | | this study | https://benchling.com/s/seq-CwGlEm2qwVg6M8Cq2fVL |
| pMNATZA1-iRFP | | this study | https://benchling.com/s/seq-ognHEdGiwTv53g9u6Izg |
| pMNATZA1-miRFP670 | | this study | https://benchling.com/s/seq-TMYFXlJAgrnfJMU2sspa |
| pMNATZA1-miRFP703 | | this study | https://benchling.com/s/seq-KF8bwpeSBJsn4IAVpCRh |
| pMNATZA1-iRFP-spmNeonGreen | | this study | https://benchling.com/s/seq-LUTpnIp22MncJeIxnja8 |

**Table S2.** ***Schizosaccharomyces pombe*** **strain list**

| Strain name | Genotype | Fig. | Source |
|---|---|---|---|
| L972 | h- | Fig. 2C, 2D, Fig. S2C, S2D | NBRP |
| L975 | h+ | | NBRP |
| SK276 | h- 2L::Padh1-NLS-iRFP-NLS<<bsd | Fig. 1B, 1C, 1D, Fig. 2B, 2C, 2D, Fig. 3B, 3C, 3G, Fig. S6A, S6C, Fig. S8B, S8C | this study, L972 |
| SK277 | h+ 2L::Padh1-NLS-iRFP-NLS<<bsd | | this study, L975 |
| SK284 | h- 2L::Padh1-NLS-iRFP-NLS<<bsd 1L::Padh1-HO1<<nat | Fig. 2B, Fig. 3B, 3C, 3G, Fig. S6B, S6D | this study, SK276 |
| SK285 | h+ 2L::Padh1-NLS-iRFP-NLS<<bsd 3R::Padh1-btFnr-bFd<<nat | Fig. 2B | this study, SK277 |
| SK287 | h+ 2L::Padh1-NLS-iRFP-NLS<<bsd z::Padh1-PcyA<<nat | Fig. 2B | this study, SK277 |
| SK294 | h- 2L::Padh1-NLS-iRFP-NLS<<bsd 1L::Padh1-HO1<<nat 3R::Padh1-btFnr-bFd<<kan | Fig. 2B | this study, SK284 |
| SK296 | h+ 2L::Padh1-NLS-iRFP-NLS<<bsd z::Padh1-PcyA<<nat 1L::Padh1-HO1<<kan | Fig. 2B | this study, SK287 |
| SK295 | h+ 2L::Padh1-NLS-iRFP-NLS<<bsd z::Padh1-PcyA<<nat 3R::Padh1-btFnr-bFd<<kan | Fig. 2B | this study, SK287 |
| SK305 | h+ 2L::Padh1-NLS-iRFP-NLS<<bsd 1L::Padh1-HO1<<nat 3R::Padh1-btFnr-bFd<<kan z::Padh1-PcyA<<nat | Fig. 2B, Fig. 3G, Fig. S6B, S6D | this study, SK294 x SK295 |
| SK282 | h- 2L::Padh1-NLS-iRFP-NLS<<bsd 1L::Padh1-SynPCB2.1<<kan | Fig. 3G, Fig. S6B, S6D, Fig. S8B, S8C | this study, SK276 |
| YG658 | h- 1L::Padh1-SynPCB2.1<<bsd | | this study, L972 |
| YG1074 | h- 1L::Padh1-SynPCB2.1<<bsd cdc2-iRFP<<kan | Fig. 4B | this study, YG658 |
| YG1096 | h- 1L::Padh1-SynPCB2.1<<bsd rpb9-iRFP<<kan | Fig. 4B | this study, YG658 |
| YG1095 | h- 1L::Padh1-SynPCB2.1<<bsd rpa49-iRFP<<kan | Fig. 4B | this study, YG658 |
| YG1085 | h- 1L::Padh1-SynPCB2.1<<bsd swi6-iRFP<<kan | Fig. 4B | this study, YG658 |
| YG1093 | h- 1L::Padh1-SynPCB2.1<<bsd pds5-iRFP<<kan | Fig. 4B | this study, YG658 |
| YG1083 | h- 1L::Padh1-SynPCB2.1<<bsd cut11-iRFP<<kan | Fig. 4B | this study, YG658 |
| YG1094 | h- 1L::Padh1-SynPCB2.1<<bsd mal3-iRFP<<kan | Fig. 4B | this study, YG658 |
| YG1084 | h- 1L::Padh1-SynPCB2.1<<bsd sfi1-iRFP<<kan | Fig. 4B | this study, YG658 |

| | | | |
|---|---|---|---|
| YG1086 | h- 1L::Padh1-SynPCB2.1<<bsd cox4-iRFP<<kan | Fig. 4B | this study, YG658 |
| YG1092 | h- 1L::Padh1-SynPCB2.1<<bsd cnx1-iRFP<<kan | Fig. 4B | this study, YG658 |
| SK323 | h- 1L::Padh1-SynPCB2.1-Lifeact-iRFP<<bsd | Fig. 5A | this study, L972 |
| YG1082 | h- 1L::Padh1-synPCB2.1-Tadh1-Padh1-NLS-iRFP-NLS-Tadh1<<bsd | Fig. 5B | this study, L972 |
| SK356 | h- 3R::Padh1-spnTurquoise2-GL-ras1delN200<<ble | | this study, L972 |
| YG1114 | h- 3R::Padh1-spnTurquoise2-GL-ras1delN200<<ble mis12-spmNeonGreen<<kan 1L::Padh1-synPCB2.1-Lifeact-iRFP<<bsd | | this study, SK356 |
| YG1124 | h- 3R::Padh1-spnTurquoise2-GL-ras1delN200<<ble mis12-spmNeonGreen<<kan 1L::Padh1-synPCB2.1-Lifeact-iRFP<<bsd z::Padh15-mCherry-atb2<<nat | | this study, YG1114 |
| YG1127 | h- 3R::Padh1-Turquoise2-GL-ras1delN200<<ble mis12-spmNeonGreen<<kan 1L::Padh1-synPCB2.1-Lifeact-iRFP<<bsd z::Padh15-mCherry-atb2<<nat c::Padh11-NLS-mTagBFP2-NLS<<hyg | Fig. 5C | this study, YG1124 |
| SK064 | h- 1L::Padh1<<nat | Fig. S2C, S2D | this study, L972 |
| SK098 | h- 2L::Padh1<<nat | Fig. S2C, S2D | this study, L972 |
| SK100 | h- 3R::Padh1<<nat | Fig. S2C, S2D | this study, L972 |
| SK062 | h- z::Padh1<<nat | Fig. S2D | this study, L972 |
| SK063 | h- z::Padh1-spmNeonGreen<<nat | Fig. S2E | this study, L972 |
| SK027 | h- 1L::Padh1-spmNeonGreen<<nat | Fig. S2E | this study, L972 |
| SK099 | h- 2L::Padh1-spmNeonGreen<<nat | Fig. S2E | this study, L972 |
| SK101 | h- 3R::Padh1-spmNeonGreen<<nat | Fig. S2E | this study, L972 |
| YG503 | h- 1L::Padh1-SynPCB2.1<<kan | Fig. S8B, S8C | this study, L972 |
| SK408 | h- 1L::Padh1-MTS-HO1<<kan | | this study, L972 |
| SK409 | h- z::Padh1-iRFP-spmNeonGreen<<nat | Fig. S4A, S4B, S4C, Fig. S5C | this study, L972 |
| SK410 | h- 1L::Padh1-synPCB2.1<<bsd z::Padh1-iRFP-spmNeonGreen<<nat | Fig. S4A, S4B, S4C, Fig. S5B, S5C | this study, YG658 |
| SK417 | h- 1L::Padh1-MTS-HO1<<kan z::Padh1-iRFP-spmNeonGreen<<nat | Fig. S4A, S4B, S4C, Fig. S5C | this study, SK408 |
| SK402 | h- z::Padh1-iRFP<<nat | Fig. S7 | this study, L972 |

| | | | |
|---|---|---|---|
| SK403 | h- z::Padh1-miRFP670<<nat | Fig. S7 | this study, L972 |
| SK404 | h- z::Padh1-miRFP703<<nat | Fig. S7 | this study, L972 |

**Table S3. Primer list**

| **Primer name** | **Primer sequence (5' → 3')** |
|---|---|
| TtefF | ATGCGAAGTTAAGTGCGCAG |
| 1Llocus-check-R | TCGGATAGTAGTTGCCAACAGC |
| 2Llocus-check-R | CGTATTTCGCTGTACATAGCATAATTTC |
| 3Rlocus-check-R | TGCAGCTGTAGTAAATTTCGAAGTC |

**Table S4. Genome information and BLASTp results**

| Species | BLASTp e-value (HO-1) | BLASTp e-value (HMX1) | Accession No. | Reference |
|---|---|---|---|---|
| Rhizopus delemar RA 99-880 | 5.19E-53 | | GCA_000149305.1 | (Ma et al., 2009) |
| Coprinopsis cinerea Okayama-7 | 1.89E-31 | 2.89E-07 | GCA_000182895.1 | (Stajich et al., 2010) |
| Laccaria bicolor S238N-H82 | 1.51E-30 | | GCA_000143565.1 | (Martin et al., 2008) |
| Ustilago maydis strain 521 | 5.92E-15 | | GCA_000328475.2 | (Kämper et al., 2006) |
| Puccinia graminis f. sp. tritici strain CRL 75-36-700-3 | | | GCA_000149925.1 | (Duplessis et al., 2011) |
| Saitoella complicata NRRL Y-17804 | 5.58E-13 | 1.03E-27 | GCA_001661265.1 | (Riley et al., 2016) |
| Neolecta irregularis DAH-3 | | | GCA_001929475.1 | (Nguyen et al., 2017) |
| Taphrina deformans PYCC 5710 | 4.16E-08 | 9.95E-24 | GCA_000312925.2 | (Cissé et al., 2013) |
| Schizosaccharomyces japonicus yFS275 | | | GCA_000149845.2 | (Rhind et al., 2011) |
| Schizosaccharomyces pombe L972 | | | GCA_000002945.2 | (Wood et al., 2002) |
| Yarrowia lipolytica | 6.73E-10 | 7.60E-63 | GCA_000002525.1 | (Dujon et al., 2004) |
| Candida albicans WO-1 | 9.74E-12 | 8.36E-56 | GCA_000149445.2 | (Butler et al., 2009) |
| Saccharomyces cerevisiae S288C | 6.52E-09 | 0 | GCA_000146045.2 | (Goffeau et al., 1996) |
| Tuber melanosporum Mel28 | | 1.07E-22 | GCA_000151645.1 | (Martin et al., 2010) |
| Aspergillus nidulans FGSC-A4 | | | GCA_000149205.2 | (Arnaud et al., 2012) |
| Neurospora crassa OR74A | | | GCA_000182925.2 | (Galagan et al., 2003) |
